# A nonhuman primate model mirrors human congenital cytomegalovirus infection and reveals a spectrum of vertical transmission outcomes

**DOI:** 10.1101/2025.01.16.633406

**Authors:** Tabitha Manuel, Matilda Moström, Chelsea M. Crooks, Angel Davalos, Richard Barfield, Elizabeth Scheef, Savannah Kendall, Cecily C. Midkiff, Lesli Sprehe, Macey Trexler, Francis Boquet, Monica Shroyer, Victoria Danner, Lara Doyle-Myers, Carolyn Weinbaum, Anne Mirza, Stephen Lammi, Claire Otero, Marissa R. Lee, Layne W. Rogers, Joshua Granek, Kouros Owzar, Daniel Malouli, Klaus Früh, Timothy Kowalik, Cliburn Chan, Sallie R. Permar, Robert V. Blair, Amitinder Kaur

**Affiliations:** Tulane National Primate Research Center; Tulane University, Covington, LA, USA; Department of Pediatrics, Weill Cornell Medicine; New York, NY, USA; Duke University School of Medicine; Durham, NC, USA; Duke Cancer Center Institute, Duke University School of Medicine, Durham, NC, USA; University of Massachusetts Chan Medical School; Worcester, MA, USA; Oregon National Primate Research Center, Oregon Health and Science University; Beaverton, OR, USA

## Abstract

Congenital cytomegalovirus (cCMV) is the leading infectious cause of birth defects worldwide, yet immune determinants of protection to inform design of a maternal vaccine remain elusive. Here, we characterized the outcome of primary rhesus CMV (RhCMV) infection during pregnancy in an immune competent nonhuman primate (NHP) model. RhCMV DNA was detected in amniotic fluid and/or fetal tissues in six of 12 (50% placental transmission) CMV-naive rhesus macaque dams inoculated intravenously with RhCMV in early second trimester gestation. Widespread tissue dissemination dominated by one of two inoculated RhCMV strains was present in one fetus (8.3% cCMV disease). Placental RhCMV transmission was associated with elevated fetal and maternal plasma TNF-alpha and reduced maternal brain-derived neurotrophic factor and IL-10 levels. CMV exposure during pregnancy had a broad impact on the placenta and fetus even in the absence of congenital infection as evidenced by RhCMV infection at the maternal-fetal interface in all 12 dams, along with significantly reduced placental efficiency and fetal growth metrics compared to gestation-matched control pregnancies. This NHP model recapitulates key aspects of human cCMV and provides new insight into barriers and biomarkers of successful vertical transmission.

**One sentence summary:** The nonhuman primate model mirrors the epidemiology of human congenital CMV (cCMV) after primary infection and reveals its transmission bottlenecks.

## INTRODUCTION

Congenital cytomegalovirus (cCMV), the leading cause of *in utero* infections, impacts approximately 1 in 200 births each year worldwide (1, 2). Following intrauterine transmission, cCMV can cause teratogenic effects such as microcephaly, intrauterine growth restriction (IUGR), developmental delays, and sensorineural hearing loss (SNHL) (2–4). Despite its global impact, there are currently no available efficacious preventatives for cCMV.

One barrier to developing a successful maternal CMV vaccine is our limited understanding of immune determinants necessary for preventing intrauterine transmission (5, 6). Primary infection of CMV-seronegative women during pregnancy has a higher risk of intrauterine transmission (30-70%) compared to reinfection or reactivation in CMV-seropositive women (1-4% transmission risk), suggesting pre-existing maternal immunity can partially protect against cCMV (5, 7). Yet it is well-established that a robust CMV-specific humoral and cellular immune response induced by natural infection is not sufficient to prevent reinfection or cCMV (8) (9) (10). Protection may require adequate immunity at the tissue level, in this instance at the maternal-fetal (M-F) interface to prevent CMV from crossing the placental barrier and infecting the fetus. Tissue immunity is difficult to assess in clinical settings due to ethical limitations and limited sample availability, so establishing a translational animal model of cCMV is critical for better defining immune determinants of protection.

Rhesus macaques have been used for studies on human CMV (HCMV) pathogenesis in the context of AIDS and transplant settings (11). This model is also ideal for studying the maternal immune system and outcomes of cCMV infection because of biologic and genetic similarities between rhesus CMV (RhCMV) and HCMV, and the physiologic and immunologic similarities between macaque and human placenta (12–14). Historical studies showed intraperitoneal, intracranial, and intraamniotic RhCMV inoculation in rhesus macaques results in fetal pathology consistent with human cCMV infection (11, 15). More recently, we established the placental transmission model of cCMV in rhesus macaques and demonstrated the ability of RhCMV to cross the placenta and cause fetal infection after intravenous inoculation in RhCMV-seronegative rhesus macaques that were either immunocompetent or CD4+ T cell-depleted prior to RhCMV infection. In the CD4+ T cell-depleted group, six of six dams (100%) transmitted RhCMV across the placenta and five (83%) experienced spontaneous abortion (16, 17). In the immunocompetent group, two of three dams transmitted RhCMV across the placenta, but did not exhibit RhCMV-associated fetal sequelae (16). Due to the small size of the immunocompetent cohort, we were unable to fully characterize intrauterine transmission in the setting of an intact immune system. We therefore set out to define a robust translational model of primary cCMV transmission using a larger immunocompetent cohort of rhesus macaques.

Here, we report the outcome of primary infection in 12 immunocompetent rhesus macaque dams inoculated intravenously with two RhCMV strains during early second trimester gestation and show that vertical transmission and cCMV disease rates in this model are like primary HCMV infection. Using whole-genome deep sequencing, we characterized the dominant RhCMV strain crossing the placenta in one fetus with extensive cCMV disease. We also define virologic and pathologic responses to primary RhCMV infection in pregnancy and show its impact on the placenta and fetus even in the absence of congenital infection. Finally, we identified several candidate biomarkers of RhCMV vertical transmission in our model. Our results corroborate published evidence that rhesus macaques recapitulate human cCMV pathogenesis and provide novel insights into the complexity of primary cCMV transmission.

## RESULTS

### Subhead 1: Viremia and Viral Shedding Kinetics

To model primary human cCMV infection, we enrolled 12 pregnant RhCMV-seronegative rhesus macaque dams that were intravenously inoculated at late first trimester / early second trimester gestation with equal proportions of RhCMV strain UCD52 and a bacterial artificial chromosome-clone (BAC) with repaired full length (FL)-RhCMV and monitored RhCMV DNAemia in maternal blood, urine, saliva and amniotic fluid obtained by ultrasound-guided amniocentesis until elective C-section near term gestation, at which point placenta and fetal tissues were harvested for further analyses (**Fig. 1A, Suppl Table 1**).

**Fig. 1.**
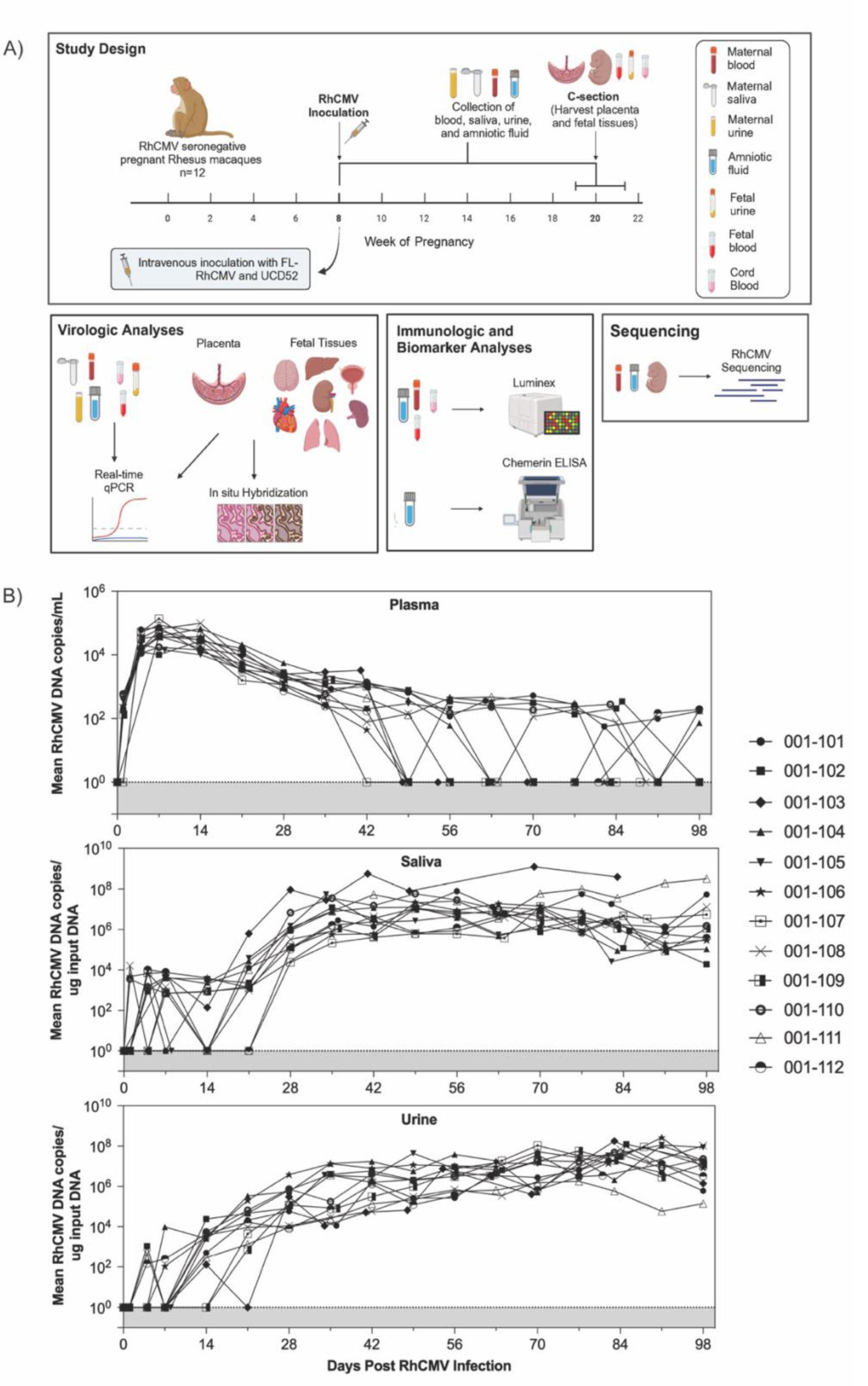
Study Design and RhCMV DNA Loads. **A)**Twelve RhCMV-seronegative, immunocompetent rhesus macaques were intravenously inoculated with two RhCMV strains during early second trimester. Blood, amniotic fluid (AF), urine, and saliva were collected weekly until elective C-section. Fetal and placental tissues were harvested at C-section. These samples were used in downstream analyses. **B)** RhCMV DNA copy numbers in plasma, saliva, and urine in individual animals. Mean of three to six technical PCR replicates for each data point shown. Viral loads expressed as mean copies of input RhCMV DNA per mL of plasma, and per ug of input DNA in saliva or urine.

RhCMV DNAemia was detected in all dams by day one post-infection (PI), peaked at day seven through 14, and thereafter began resolving with variable kinetics (**Fig. 1B and S1**). Beginning at six weeks PI, eight of 12 (75%) dams became intermittently aviremic. Viral shedding in the saliva and urine appeared within the first three weeks PI with variable kinetics among the dams. By week four PI, all dams were consistently shedding RhCMV in saliva and urine at levels ranging between 10^6^ and 10^8^ RhCMV DNA copies per ug input DNA that persisted for the duration of the study (**Fig. 1B and S1**). This is consistent with prior studies showing immunocompetent rhesus macaques become aviremic, yet persistently shed virus in saliva and urine (18).

### Subhead 2: RhCMV Intrauterine Transmission

In clinical settings, the “gold standard” method for detection of cCMV includes amniocentesis followed by PCR analysis of the amniotic fluid (AF) (19) (20). We replicated this method in our previous studies, as well as in the current study. Consistent with our prior study, AF transmission was defined by PCR detection of RhCMV DNA in at least two of 12 individual replicates of AF per time-point (21). Based on this criterion, we determined five dams were amniotic fluid positive (AF+), and the remaining seven dams were amniotic fluid negative (AF–) (**Fig. 2A**). RhCMV DNA in one of 12 replicates was detected in the AF of dam 001-102; however, this did not meet our criterion for AF positivity.

**Fig. 2.**
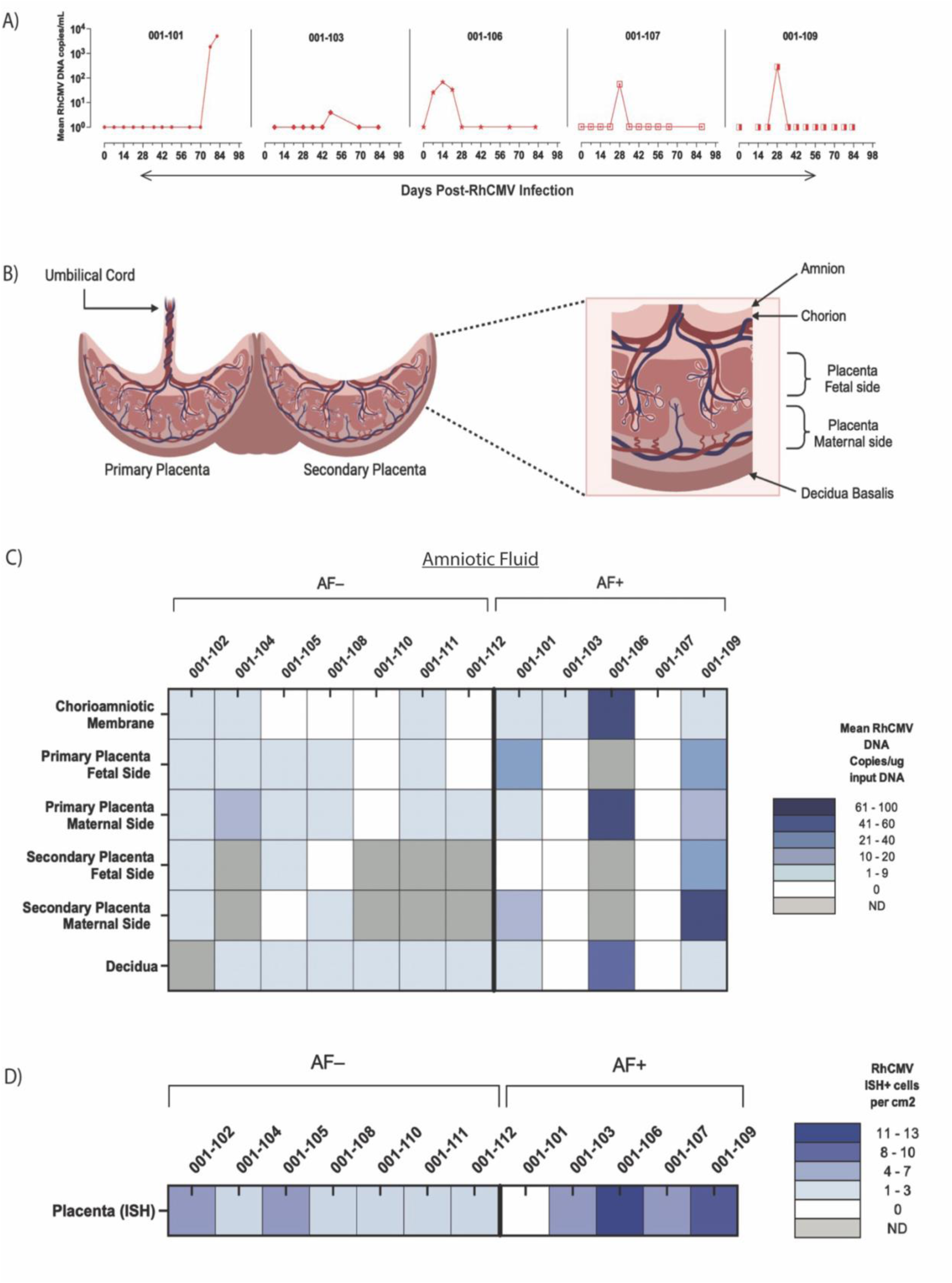
Quantification of RhCMV DNA at the Maternal-Fetal (M-F) Interface. **A)**RhCMV DNA detection in amniotic fluid (AF) of five AF+ dams. Viral loads expressed as mean RhCMV DNA copies per mL of AF. Data on mean of 12 PCR replicates per time-point. **B)** Schematic diagram of rhesus macaque M-F interface tissues. **C)** Heatmap illustrating mean RhCMV DNA copies per ug of input DNA within tissues of the M-F interface of each dam within AF+ and AF– group. Mean copy number of 12 PCR replicates shown. Gray boxes (ND) indicate no sample collected. **D)** Heatmap showing number of RhCMV-positive cells per cm^2^ by in situ hybridization (ISH) in full-thickness placental cross-sections of individual AF+ and AF– dams.

The kinetics of RhCMV DNA detection in the AF varied among the dams. It was detected transiently within seven weeks PI in four of five AF+ dams. Dam 001-106 showed the earliest detectable RhCMV DNA in the AF at three time-points (days 7, 14, and 21 PI), whereas dams 001-109, 001-107, and 001-103 had detectable RhCMV DNA at a single time-point between day 28 and day 49 PI (**Fig. 2A**). In contrast, RhCMV DNA in the AF of dam 001-101 was detected on days 77 and 82 PI and showed the highest RhCMV DNA copy numbers (**Fig. 2A**).

We also evaluated the maternal-fetal (M-F) interface for RhCMV infection. Each component of the M-F interface (amniochorion, fetal and maternal surfaces of both the primary and secondary placental discs, and maternal decidua) were dissected and RhCMV DNA quantitated by quantitative real-time PCR (**Fig. 2B-C**). *In situ* hybridization (ISH) was performed on a full thickness section (decidua through amniochorion) of the primary placental disc to confirm placental RhCMV infection (**Fig. S2A-B**). Based on our PCR positive criteria outlined in the methods, 11 of 12 dams were RhCMV DNA-positive at the M-F interface (**Fig. 2C**). 11 of 12 dams had RhCMV ISH-positive cells at the M-F interface (**Fig. 2D**). PCR and ISH quantitation of RhCMV infection at the M-F interface did not correlate and were sometimes discordant, as demonstrated by dams 001-101 and 001-107 **(Fig. 2C-D)**. We did not detect RhCMV DNA in the umbilical cord of any dam.

Taken together, all 12 dams met our criteria for being RhCMV-positive at the M-F interface. Discordance between quantitative PCR and ISH data is likely due to differences in sampling and the random multifocal distribution of RhCMV throughout the placenta. Our PCR and ISH results paint a complex picture that highlights the high frequency of detection of RhCMV at the M-F interface even in the absence of evidence of AF positivity and emphasizes the need for multimodal diagnostics to assess placental transmission risk.

### Subhead 3: cCMV Outcomes

All placental lesions identified were of mild severity. Lesions included acute to subacute placentitis in four dams; chorioamnionitis in one dam; and placental infarction in two dams (**Suppl Table 2**). Additional histopathologic changes in placentas included scattered placental fibrinoid, mineralization, and syncytial knot formation. However, these changes can be seen with variable frequency in normal, late term placentas, and therefore were not interpreted as pathologic.

One of 12 fetuses (fetus of dam 001-101) had widespread RhCMV infection based on multiple tissues with detectable RhCMV DNA by PCR (**Fig. 3A**). The highest RhCMV DNA copy numbers within this fetus were found in the parietal cortex, basal ganglion, and lung, with lower copy numbers in the cochlea, kidney, spleen, liver, submandibular salivary gland, parotid salivary gland, hippocampus, cerebellum, frontal cortex, occipital cortex, thalamus, and hypothalamus (**Fig. 3A**). RhCMV DNA was also detected in the plasma and cord blood of this fetus, but not in the urine which was collected by cystocentesis (**Fig. 3A**).

**Fig. 3.**
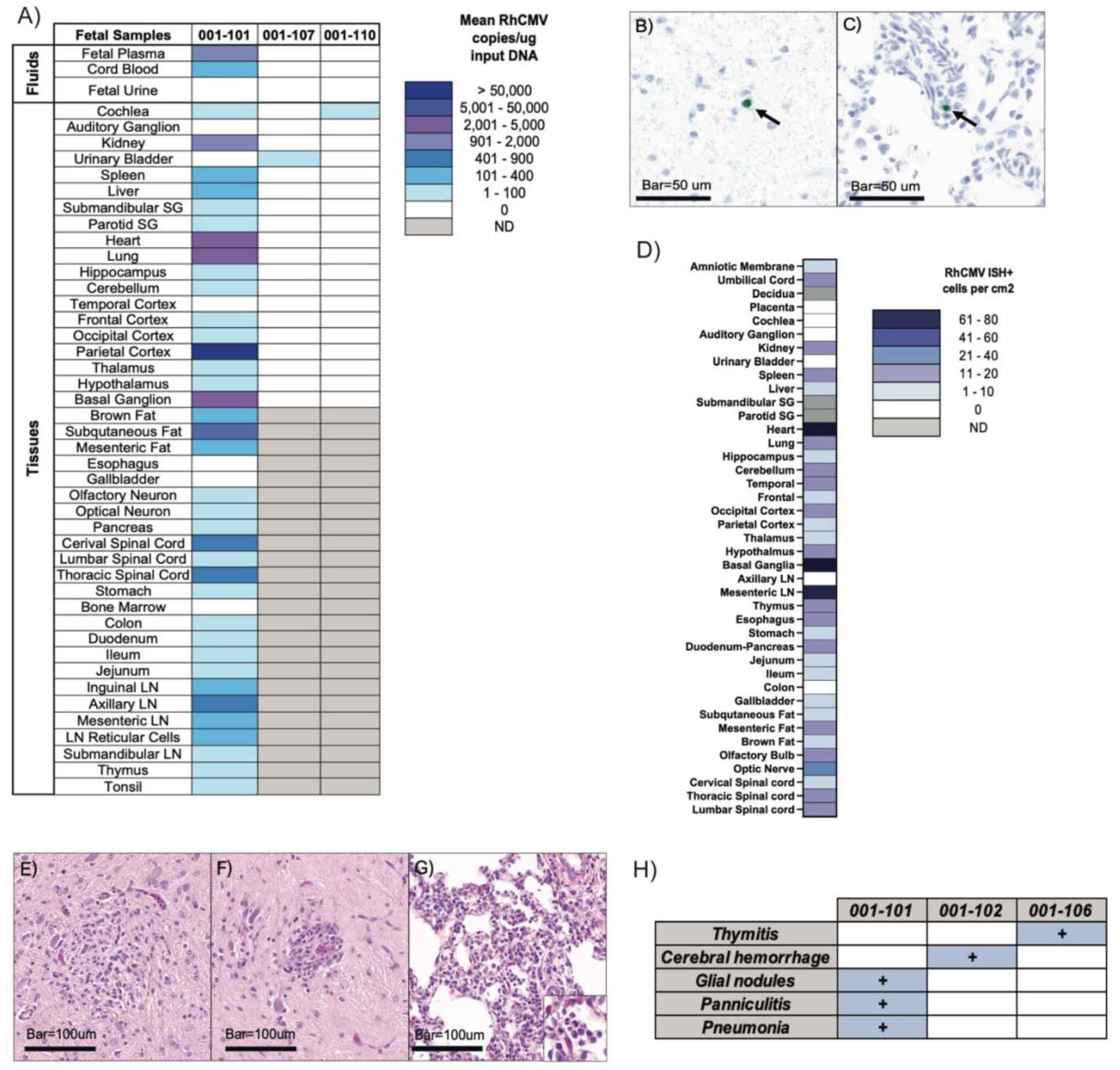
Fetal cCMV Infection and Pathology. **A)**RhCMV DNA copies in fetal tissues of dams 001-101, 001-107, and 001-110. Mean copy number of 12 PCR replicates shown and expressed per ug of input DNA. Gray boxes indicate no sample collected. **B–C)** Representative RhCMV ISH of fetal tissues from dam 001-101. Rare RhCMV RNA positive cells (green, arrows) were detected in inflamed sections of hypothalamus **(B)** and lung **(C)** with RNAscope. Bar=50 um. **D)** Heatmap showing number of RhCMV-positive cells per fetal tissue sample from dam 001-101. **E-G)** Histopathology of fetal tissues from dam 001-101 showing **E)** Glial nodules in brainstem; **F)** Hypothalamus, fetal lesions included cerebral vessels with mononuclear perivascular cuffs and scattered glial nodules (not shown); **G)** Lung, alveolar septa were thickened by inflammatory cells with rafts of neutrophils (inset) within alveoli. Bar=100um. **H)** Listing of lesions discovered during histopathological analysis of fetal tissues from dams 001-102, 001-106, and 001-101.

Two additional fetuses were RhCMV DNA positive in one organ (**Fig. 3A**). The fetus of dam 001-107 had two of 12 positive replicates in the fetal urinary bladder while the fetus of dam 001-110 had two of six positive replicates in the cochlea. Four other fetuses (001-102, 001-104, 001-105, and 001-109) had detectable levels of RhCMV DNA in one of 12 replicates in a single organ, but did not meet our criterion for PCR positivity. These included the fetal urinary bladder of 001-104, the fetal cochlea of 001-102, the parietal cortex of 001-109, and the auditory ganglion of 001-105.

To confirm the widespread infection found in the 001-101 fetus, we performed ISH on the same tissues used in our PCR analysis (**Fig. 3B-D**). Similar to the analyses of the M-F interface, RhCMV quantitation in fetal tissues by ISH and PCR did not correlate (**Fig. S2C**). However, both methods indicated widespread RhCMV infection. Viral intranuclear inclusions, which are characteristic of CMV infection, were not observed, but the presence of RhCMV was confirmed by ISH in lesions found during histopathological examination (**Fig. 3B-D**). Lesions consistent with cCMV were identified in multiple organs of the fetus of dam 001-101 (**Fig. 3E-H**) and included mild lymphocytic perivascular cuffing and multifocal glial nodules within the hypothalamus and brainstem, interstitial pneumonia, and rare panniculitis. Two other fetuses had histopathologic lesions-the fetus of dam 001-102 had a focus of cerebral hemorrhage in the frontal lobe, and the fetus of dam 001-106 had mild histiocytic infiltration of the thymus (**Fig. 3H**).

It is generally thought that the source of CMV in the AF is fetal urine, so AF positivity is considered an indicator of fetal infection. We were able to collect urine by cystocentesis from seven of 12 fetuses at fetal necropsy including four AF+ dams (001-101, 001-103, 001-107, 001-106) and three AF– dams (001-110, 001-105, 001-108). All fetal urine samples were RhCMV-negative by PCR. Fetal plasma IgM serology performed on IgG-depleted samples was also inconclusive of fetal infection (**Fig. S3A-B**).

Compared to our previous studies of primary RhCMV infection in the setting of CD4+ T lymphocyte depletion (16, 17), all dams in this cohort had reduced risk of vertical transmission (41.6% AF+ vs 100% AF+ in CD4-depleted dams) and carried to elective C-section with no incidence of spontaneous abortion (100% vs 16.7% fetal survival in CD4-depleted dams) (**Fig. S4A-B**).

### Subhead 4: Fetal and Placental Morphometrics

cCMV can cause delayed fetal growth and intrauterine growth restriction (IUGR). Therefore, we performed weekly ultrasound measurements of the fetal hand, foot, humerus, femur, occipito-frontal diameter (OFD), biparietal diameter (BPD), head circumference, abdominal circumference, and chest circumference. We limited our statistical analyses to the BPD and femur length, as these are the most reliable and clinically relevant measurements, and compared them to reference sonographic values across gestation age in normal pregnant rhesus macaques (22). No statistically significant differences were noted in the BPD and femur growth rate compared to the reference. Minor variations were noted among fetuses in our cohort between gestation days 108-136, with some of the fetuses exhibiting a shorter average femur length, and in one fetus, 001-111, a narrower BPD, but these differences were not significant **(Fig. 4A)**. There were no statistically significant differences between the fetal growth rates of AF+ and AF– dams.

**Fig. 4.**
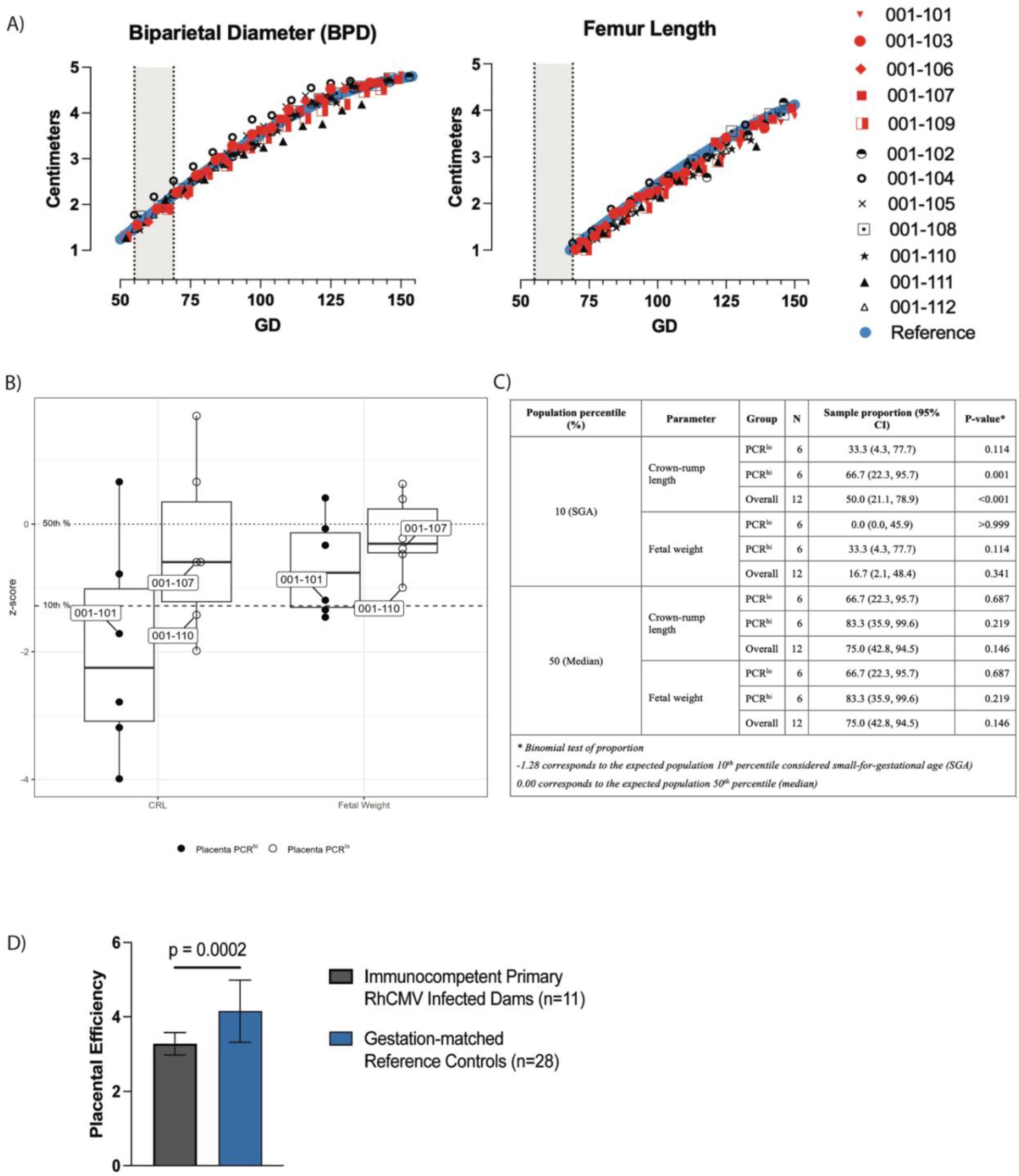
Fetal Morphometrics. **A)**Fetal growth curves of biparietal diameter (BPD) and femur length. AF+ dams denoted by red icons, AF– dams shown in black, and referent population indicated by blue icons. Shaded region and dotted line indicate range of gestation days that dams were inoculated with RhCMV. **B)** Box and whisker plot demonstrating CRL and fetal weight z-scores of placenta PCR^hi^ and PCR^lo^ dams. **C)** Summary of morphometric parameter z-scores below population median and small for gestational age (SGA) by placenta PCRhi/lo burden and overall. Bar graph illustrating placental efficiency ratios of the study dams in gray and the referent population from Roberts et al (*23*) in blue. Significance determined by Wilcoxon rank sum test.

At C-section, the fetus and placenta were weighed, and fetal measurements including crown-rump length (CRL) were taken and compared to published reference values for fetal morphometrics based on gestation day (23) (**Suppl Table 3A**). All parameters were centered and scaled by gestational age specific reference mean and standard deviation, respectively, to define z-scores based on the published values. We then assessed the z-score characteristics of CRL and fetal weight (**Fig. 4B-C**). Six of 12 fetuses (50%) had CRL z-scores values below the expected 10^th^ percentile (i.e z-score < −1.28) considered small for gestation age (SGA). Two of 12 (16.7%) fetuses had fetal weight z-scores in the SGA range (**Fig. 4B**). Our cohort had a discernably greater proportion of fetuses with SGA CRLs than expected (p <0.001; **Fig. 4C**). For CRL and fetal weight, nine out of 12 (75%) fetus z-scores were below zero (**Fig. 4C**). The observation that two-thirds of the cohort had z-scores below zero could suggest potential IUGR in primary RhCMV-infected dams.

Next, we grouped fetuses based on PCR results to determine whether detectable RhCMV DNA in the AF (n=7 AF– *vs* n=5 AF+), or fetal tissue (n=9 Fetus PCR– *vs* n=3 Fetus PCR+) correlated with fetal morphometrics measured at C-section (**Suppl Table 3A**). Additionally, we grouped fetuses based on a scoring metric we developed for placental RhCMV spread at the M-F interface as Placental PCR^lo^ (n=6) and PCR^hi^ (n=6) (described in methods) (**Suppl Table 3A**). We did not observe significant differences in morphometrics between groups based on AF-positivity or fetal infection (**Suppl Table 3B**). However, the CRL and fetal weight parameters showed a consistent separation of z-scores between the Placental PCR^lo^ and PCR^hi^ groups (**Suppl Table 3A**). The PCR^hi^ dams displayed a smaller CRL compared to the PCR^lo^ group (p=0.092; **Suppl Table 3B**). The proportion of fetuses with CRL z-scores below the 10^th^ percentile was higher in the Placenta PCR^hi^ (66.7%) compared to the Placenta PCR^lo^ (33.3%) group. This suggests a potential association between SGA and RhCMV burden at the M-F interface (**Fig. 4B-C**).

Placental efficiency (PE) is a measure of how well the placenta supports fetal nutrition, growth, waste management, and oxygen transfer. PE, thus, is an important measurement to consider when drawing conclusions about fetal and placental health following primary RhCMV infection. For each dam, we collected the weight in grams of the fetus and placenta at C-section. Total placental weight includes primary and secondary placental discs, as well as the umbilical cord. We then calculated the fetal:placental weight ratio to use as a proxy of PE (23). We compared the PE of our dams to that of similar gestational age controls from Roberts et al (23). The fetus of dam 001-109 was not weighed, so this dam’s PE was unable to be calculated. The PE of dams in our cohort (n=11) ranged from 2.71 to 3.89 (median=3.33). Comparatively, the gestation-comparable controls (n=28) taken from Roberts *et al*. had PE measurements ranging from 1.9 to 5.6 (median=4.11). Exact Wilcoxon rank sum test showed statistically significantly lower PE measurements in our cohort compared to these controls (**Fig. 4D**; p=0.0002).

### Subhead 5: Viral Sequencing

We used whole genome sequencing to understand *in vivo* dynamics of the two inoculating strains over the course of infection in 001-101, the only dam to exhibit widespread fetal infection. We performed amplicon-based Illumina deep sequencing of RhCMV from maternal plasma during acute infection (4, 7, 14 days PI), plasma and AF at the timepoints when AF was positive for RhCMV (77 and 82 days PI), and five fetal tissues at the time of delivery that had high viral loads (82 days PI; heart, lung, kidney, parietal cortex, basal ganglion). Three samples (plasma 77 and 82 days PI, and AF 82 days PI) had fewer than 1000 input copies and we observed a corresponding decrease in genome coverage in these samples.

We determined the inferred strain frequency in each sample by identifying genomic positions where the two reference strains differed and applied several filters to those positions to account for strain-specific mapping biases (**Fig. S5**). The total number of remaining single nucleotide variants (SNVs) used for each sample ranged from 14-862 (median 710) and are specified in **Suppl Table 4.** For each sample, we then calculated the mean proportion of aligned bases per position that had the nucleotide corresponding to either strain (number of reads with UCD52 or FL nucleotide/total reads) to estimate the relative abundance of each strain in the sample. To ensure our method was robust to the reference genome used for mapping, we performed these calculations using mapping outputs with either RhCMV-FL and RhCMV UCD52 as the mapping reference and found consistent inferred strain frequencies (within 1.5%) between the two mapping references **(Fig. 5, Fig. S6A, Suppl Table 4).** Furthermore, when performing our strain frequency analysis on the sequenced stock virus, the inferred strain frequency was >99.8% for both viruses **(Suppl Table 4, Fig. S5E-F).** Across all samples, mean inferred strain frequency for “other strain” was <0.03%, highlighting a low level of other variants resulting from de novo mutations or sequencing error.

**Fig. 5.**
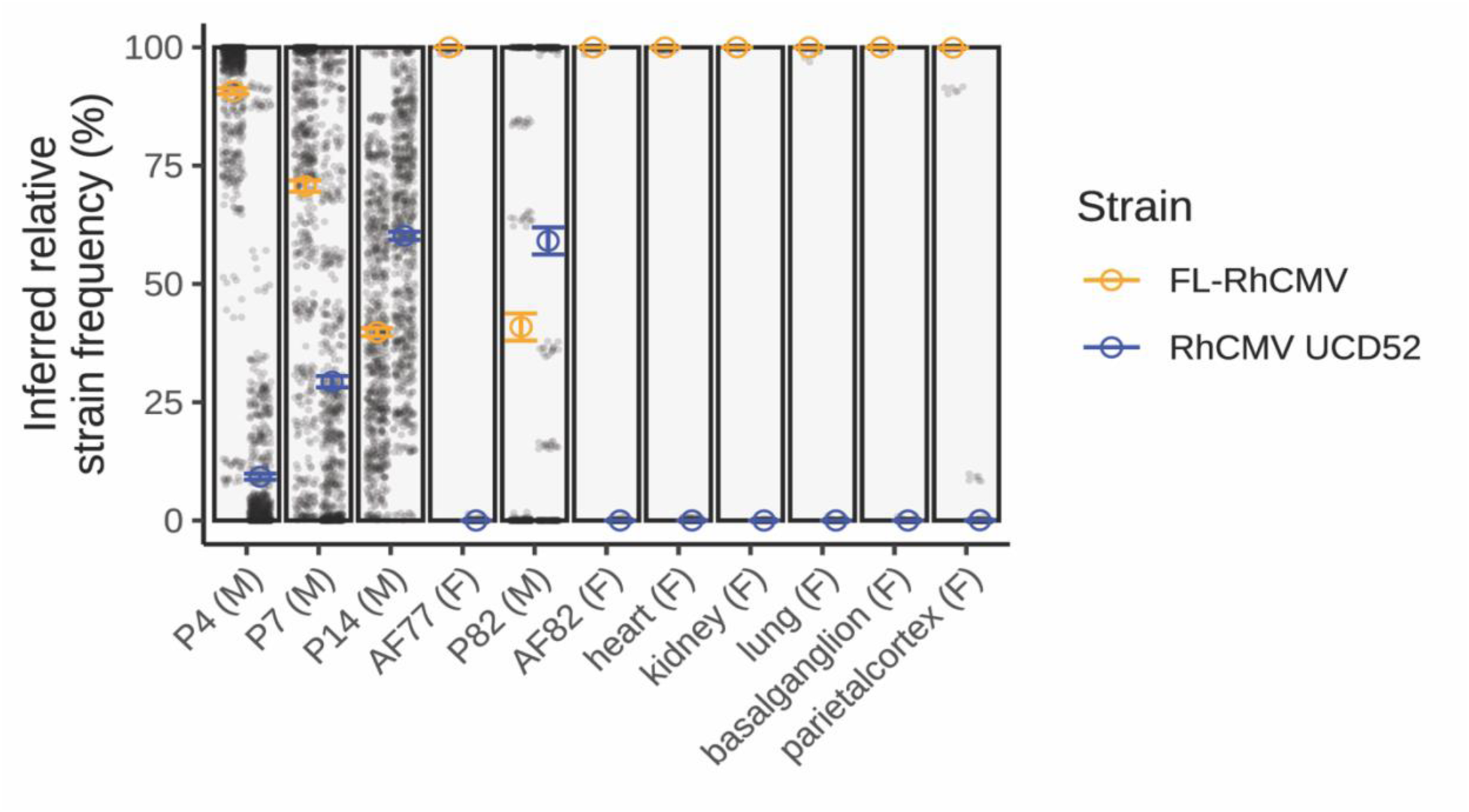
Inferred RhCMV strain frequency in animal 001-101 with FL-RhCMV as mapping reference. For each sample, inferred strain frequency of RhCMV UCD52 (blue) or FL-RhCMV (orange) was determined by taking mean of the percent of reads with the nucleotide corresponding to each strain at positions where the two inoculating strains differ. Sample IDs are denoted with “P” = plasma, “AF” = amniotic fluid, # = days post infection, “M” = maternal, and “F” = fetal. Mean indicated by open circle with error bars corresponding to standard error. Black dots represent frequencies at a single SNV.

At the acute plasma timepoints (4, 7, 14 days PI), FL-RhCMV virus is the dominant replicating strain early in infection (90-91% at 4 days PI), but by 14 days PI, the strains appear to be present in a mixed population (39-40% FL, 60-61% UCD52; **Fig. 5**). Although we sequenced virus from plasma at 77 days PI, the overall genome coverage was low due to limited input virus, and only 14 SNVs were retained after filtering for determining inferred strain frequency (**Fig. S6, panel E**). Therefore, we omitted this sample from our inferred strain frequency calculations. AF from 77 days PI and the paired AF and plasma samples from 82 days PI, despite an apparently mixed population of virus in the plasma at 82 days PI, FL-RhCMV appears to be at nearly 100% frequency in the AF. Of note, there was limited virus present in the plasma at 82 days PI, so the inferred strain frequency is based on 273-275 SNVs. We sequenced five fetal tissues, also collected at 82 days PI, and these tissues appear to have nearly 100% FL-RhCMV, consistent with the AF (**Fig. 5**).

### Subhead 6: Acute Host Response Association with cCMV Transmission

We evaluated the early host response as a predictor of RhCMV infection outcome. White blood cell (WBC) counts revealed a transient decline in leukocytes (p=0.001; p_adj_=0.0029) by day seven PI due to a decline in neutrophil counts **(Fig. 6A**). This was followed by an increase in lymphocyte counts that persisted up to day 84 PI **(Fig. 6A**). The mean absolute monocyte counts significantly increased by the third week PI (p=0.0005; p_adj_=0.0015), then returned to baseline levels by week six PI (**Fig. 6A**).We observed no discernable differences between cell counts of AF+ and AF– dams.

**Fig. 6.**
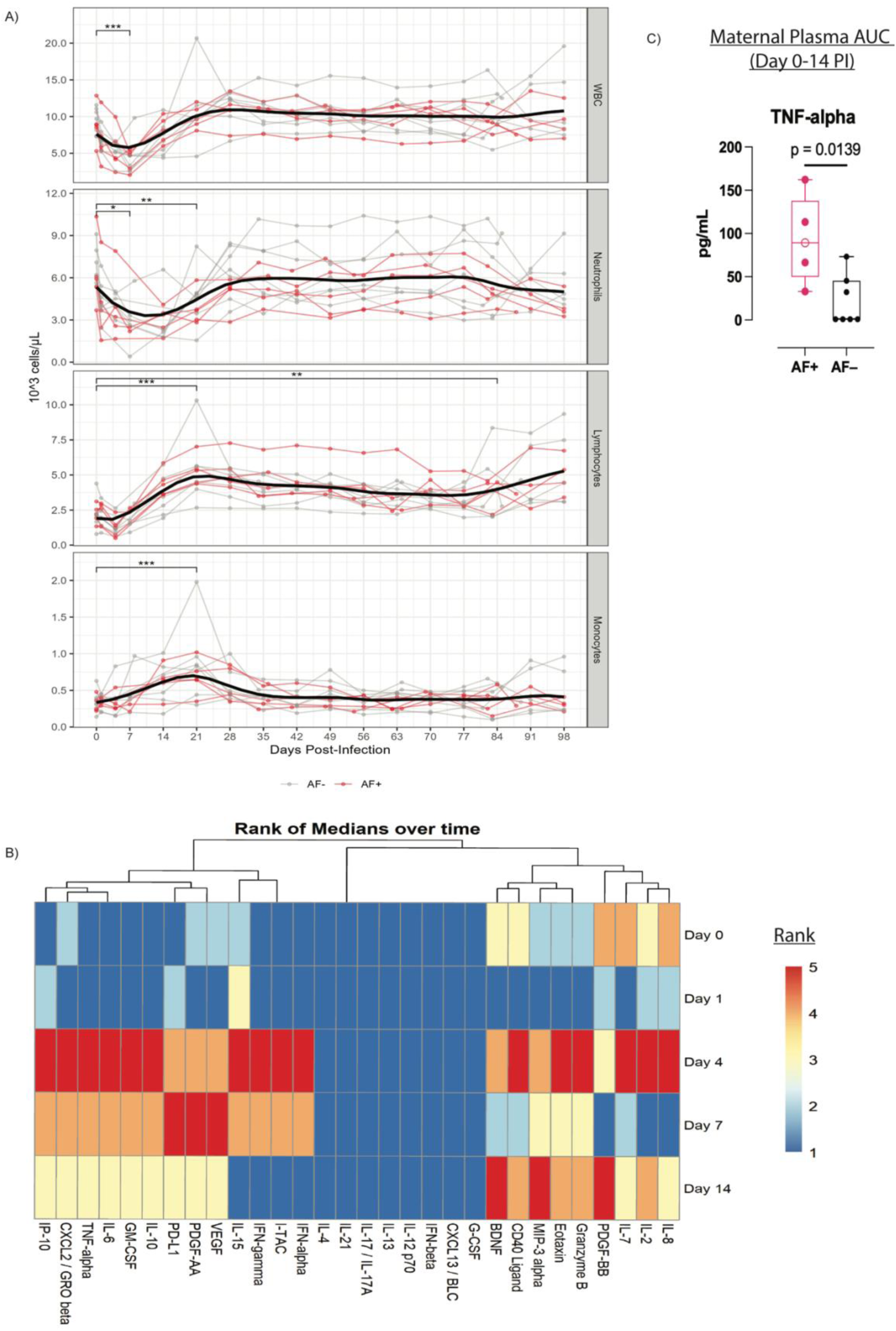
Acute Host Response to Primary RhCMV Infection. **A)**Mean absolute counts of circulating total white blood cells (WBC), neutrophils, lymphocytes, and monocytes from day 0 to day 98 post-infection (PI). Gray lines denote AF– dams, while red lines denote AF+ dams. **B)** Heatmap illustrating rank values of median maternal plasma concentrations of cytokines between 0 and 14 days PI. Cytokines measured by a 33-plex NHP Luminex assay. **C)** Box and whisker plot illustrating area under the curve (*16*) of maternal plasma TNF-alpha concentrations during first two weeks PI. Red icons denote AF+ dams and black icons refer to AF– dams. The open red circle denotes dam 001-101. Significance determined by Wilcoxon rank sum test.

Using a 33-plex NHP Luminex assay, we evaluated concentrations of growth factors, Th1 and Th2 cytokines, and several chemokines within the maternal plasma in the first two weeks PI. Of the 30 analytes which passed our quality control requirements, 16 demonstrated highest median concentrations on day four PI, three at day seven PI, and three at day 14 PI (**Fig. 6B**). Multiple analytes showed changes over time with several showing a significant post-hoc difference against baseline (**Suppl Table 5**). Eight analytes displayed no change in median concentrations across the first two weeks PI (**Fig. 6B and Suppl Table 5**). Several pro-inflammatory and anti-viral analytes showing a statistically significant transient increase at day four PI included GM-CSF, IFN-alpha, IP-10, IL-6, and VEGF (**Suppl. Table 5**). When comparing the area under the curve of analyte concentrations in maternal plasma during the first two weeks post-infection, one analyte, TNF-alpha, was higher in AF+ dams compared to AF– dams (p=0.0139; p_adj_: 0.417 **Fig. 6C**).

### Subhead 7: Candidate Biomarkers of Primary cCMV Transmission

PCR analyses showed a significant increase in cumulative viral load (via AUC) within the urine of AF– dams compared to AF+ dams during the first two weeks PI (p= 0.0177, p_adj_: 0.053, **Fig. S7A**). There was no association between AF status and the AUC of viral loads in plasma or saliva during the study. When we examined cumulative viral load over the remainder of the study, we no longer observed the statistically significant association in urine (**Fig. S7B and S7C**).

We then evaluated analytes within maternal plasma, AF, and fetal plasma collected at C-section for differences between AF+ and AF– dams. In the maternal plasma, AF+ dams had lower concentrations of brain-derived neurotrophic factor (BDNF) (p=0.048, p_adj_: 0.502) and IL-10 (p=0.0455, p_adj_: 0.502) compared to AF– dams (**Fig. 7A and Suppl Table 6**). In contrast, IFN-alpha (p=0.0265; p_adj_: 0.342) and IL-21 (p=0.0467; p_adj_: 0.342) were higher in the AF of AF+ dams (**Fig. 7B and Suppl Table 7**). Remarkably, fetal plasma TNF-alpha concentrations were significantly higher in AF+ dams compared to AF– dams, and this remained statistically significant after multiple testing correction (**Fig. 7C**; p=0.00126; p_adj_: 0.0379; **Suppl Table 8**). This difference was not observed in cord blood plasma **(Suppl Table 9)**.

**Fig. 7.**
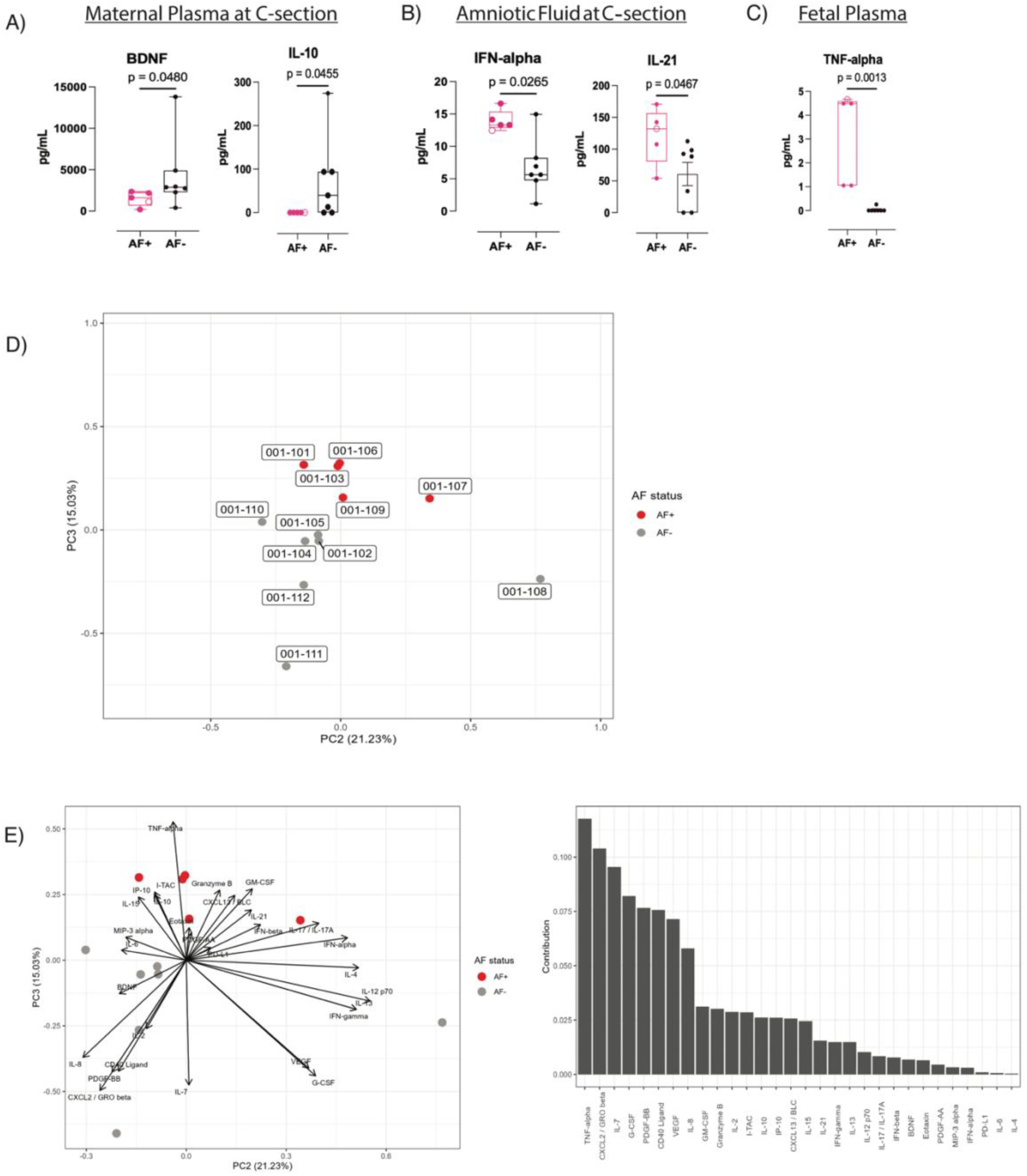
Evaluation of Biomarkers of cCMV Transmission. **A)**Concentrations of BDNF and IL-10 in maternal plasma at time of C-section for AF+ dams (red) and AF– dams (black). **B)** Plots showing concentrations of IFN-alpha and IL-21 in the amniotic fluid at C-section in AF+ dams (red) and AF– dams (black). **C)** Plot displaying TNF-alpha concentrations in fetal plasma of AF+ dams (red) and AF– dams (black). Open circle in A-C denotes dam 001-101. **D)** Principal Components Analysis (PCA) visualization shows clear separation of AF+ (red) and AF– dams (gray) in PC3 axis when evaluating analytes in fetal plasma. **E)** Biplot (left) reflecting importance of each variable with regards to the principal components. Bar graph (right) illustrates the importance of each analyte in regard to the principal components. Significance determined by Wilcoxon rank sum test.

While not statistically significant, other notable findings included higher concentrations of IFN-beta, IFN-gamma, and VEGF (p=0.0732; p_adj_: 0.342) in the AF of AF+ dams (**Suppl Table 7**). As in the maternal plasma at C-section, levels of BDNF in the AF were lower in AF+ dams compared to AF– dams (p=0.0732; p_adj_: 0.342; **Suppl Table 7**). We observed higher granzyme B concentrations in cord blood of AF+ dams compared to AF– dams (p=0.0568; p_adj_: 0.795; **Suppl Table 9**).

We then performed exploratory principal component analysis (PCA) on maternal plasma, cord blood, fetal plasma, and AF cytokine data. Only the PCA of fetal plasma showed separate clusters for AF+ and AF– dams evident in the PC3 axis (**Fig. 7D**). This difference was primarily driven by TNF-alpha, followed by IL-7 and CXCL2 (**Fig. 7E**).

Vorontsov *et al.* identified the protein chemerin levels in mid-gestation AF as a prognostic biomarker of severe cCMV infection in humans (24). We used AF samples obtained during gestation weeks seven through nine (equivalent of early second trimester in human pregnancies) to assess this analyte in our model and did not observe any difference between AF+ and AF– dams either during the second trimester (p=0.7857; **Fig. S8**) or at C-section (p=0.7879; **Fig. S8**). Dam 001-101, which displayed widespread fetal infection, did not have high levels of chemerin compared to other dams in the cohort.

## DISCUSSION

In this study we established the outcomes of primary CMV infection during pregnancy in immunocompetent rhesus macaques with a longitudinal systemic and tissue-level analysis of events leading to congenital transmission. We demonstrate that the rhesus macaque model recapitulates key features of human cCMV including the rates of vertical transmission after early second trimester primary infection, the profile of fetal tissues infected *in utero*, and detection of disseminated CMV disease in the newborn at a frequency akin to symptomatic cCMV in humans. We also report a broad outcome of maternal RhCMV exposure during pregnancy ranging from M-F interface infection and deleterious effect on fetal growth without vertical transmission to vertical transmission with disseminated fetal infection. Our results validate the translational utility of this model for human cCMV studies and broadens our understanding of bottlenecks to virus passage across the placenta.

Our cohort showed a 50% vertical transmission rate based on RhCMV DNA detection in AF (n=5) and fetal tissue without AF DNA positivity (n=1). We also identified an 8% rate of widespread fetal infection-similar to the rate at which symptomatic cCMV is reported in humans (2). Dam 001-101, which had widespread fetal infection, exhibited a distribution of virus within fetal tissues consistent with reports in human cCMV cases (25). Consistent with our previous findings of primary infection in CD4-depleted dams (17), one of the RhCMV-positive tissues was the cochlea, which also corroborates findings from murine studies that show CMV migrates to the inner ear after inoculation of the cerebral cortex (26). RhCMV localization to the cochlea may be indicative of inner ear pathology that leads to sensorineural hearing loss – a common outcome of human cCMV (26, 27). RhCMV DNA was also detected in tissues consistent with CMV natural history (urinary bladder) and organs involved in cCMV symptomatology (brain, cochlea, and auditory ganglion). The fetus of dam 001-101 exhibited encephalitis, pneumonia, and panniculitis. This is consistent with CMV-associated sequelae in humans, although our observations in this fetus were not as severe outcomes in human infants (25, 28, 29). Much of the current understanding of human cCMV pathology is limited to cases with severe outcomes, such as those with spontaneous abortion or birth defects, which did not occur in our study. This likely explains why severe pathological outcomes reported in clinical studies were not observed in our cohort (25, 29-32).

Although fetal urine is thought to be the major source of CMV in AF, other sources of viral entry into AF are also described (33–35). In our model, RhCMV PCR of fetal urine collected by cystocentesis was not an accurate diagnostic of vertical transmission. All fetal urine tested was RhCMV-negative, including fetuses of AF+ dams and the fetus of dam 001-101 which showed widespread fetal tissue infection, including infection of the kidney. Because CMV is a slow replicating virus, successful replication within the fetal kidney and subsequent excretion of virus may not occur until up to three weeks post-infection of the fetus (33). If this fetus became infected near the time of C-section, the virus may not have undergone sufficient replication for shedding in the urine. Moreover, the replicating virus may be more present in the lower urinary tract which is not fully assessed by urine collected by cystocentesis. The fetus of dam 001-101 did have detectable levels of RhCMV DNA in the lungs and salivary glands. Thus, fetal saliva and pulmonary secretions may be the alternative source of RhCMV in the AF of this dam, consistent with the current understanding of AF contents (33–35).

When using current standards for clinical diagnosis of cCMV, our model demonstrated the expected epidemiological and virologic patterns of primary cCMV transmission. However, a more in-depth interrogation of the M-F interface, AF, and fetal tissues by PCR and ISH revealed additional outcome cohorts (**Fig. 8A**). The cohorts are characterized by the following features: 1) RhCMV detection at the M-F interface without AF positivity or fetal infection, implying infection at the M-F interface does not always lead to fetal virus exposure; 2) RhCMV detection at the M-F interface with AF positivity, but negative fetal tissue, implying placental transmission has occurred without establishing infection in the fetus; 3) RhCMV at the M-F interface with AF and/or limited fetal tissue positivity, implying vertical transmission with possible asymptomatic infection of the fetus; 4) RhCMV at the M-F interface with AF positivity and widespread fetal infection, implying disseminated, symptomatic cCMV infection. These outcome cohorts paint a more complicated, non-binary picture of cCMV transmission, with the potential for unrecognized clinical impacts on the developing fetus. Of note, reports of detection of CMV DNA in late gestation human placenta from normal deliveries (36–39) suggests that non-clinically recognized CMV infection at the M-F interface may not be unique to the rhesus macaque model.

**Fig. 8.**
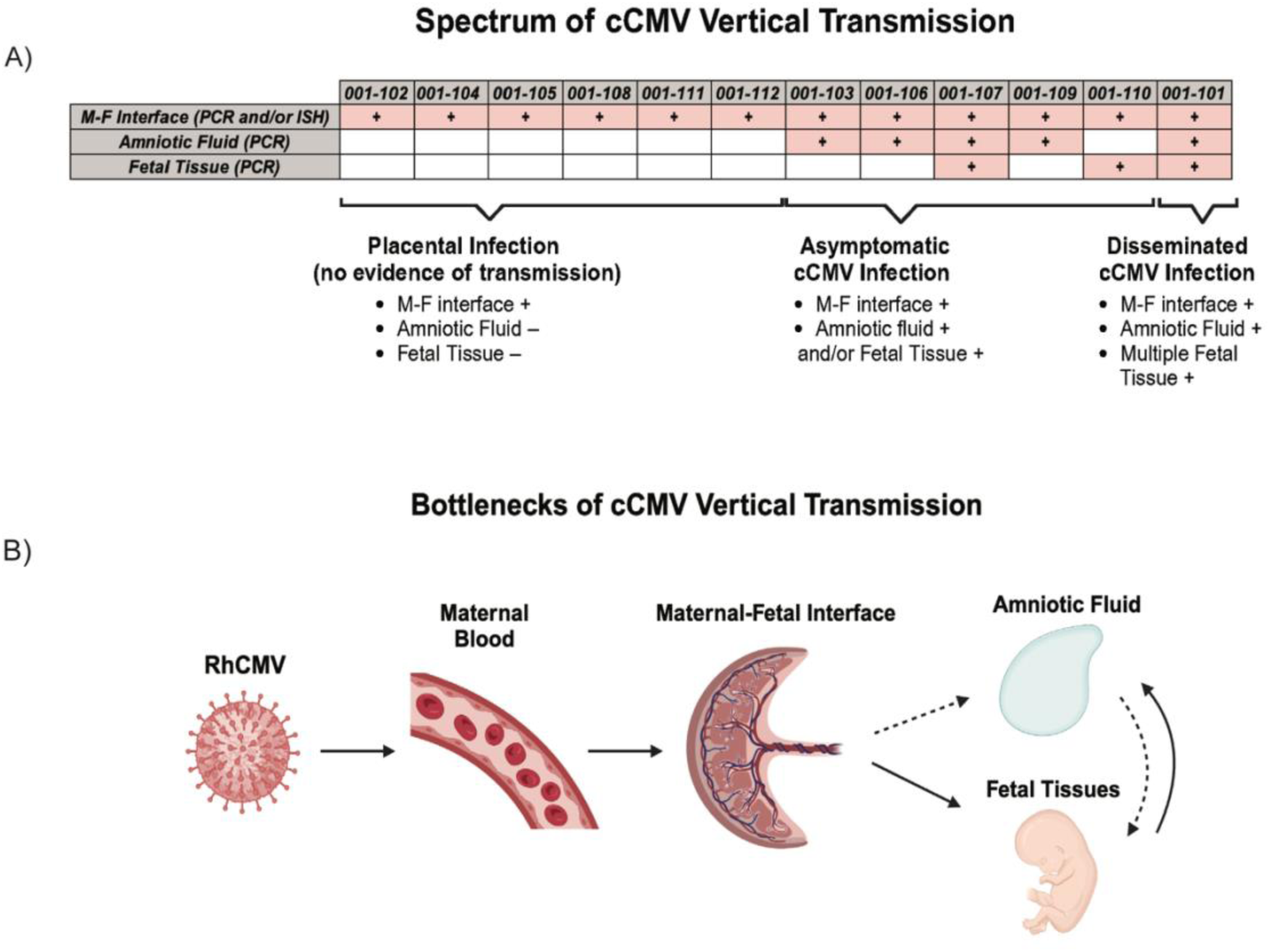
Bottlenecks of cCMV Intrauterine Transmission. **A)**Schematic summarizing PCR and ISH results in Amniotic Fluid (AF), M-F interface, and fetal tissues. Pink cell/plus sign indicate RhCMV nucleic acid detection in the respective tissues. Below schematic is a description of our proposed spectrum of congenital RhCMV intrauterine transmission. **B)** Diagram of possible routes of cCMV vertical transmission. Dotted lines represent new proposed modes of vertical transmission found in our cohort.

The route by which CMV crosses the placenta is not fully understood, but placental HCMV infection in the absence of fetal infection suggests that the placental barrier is critical for protection (40). Similarly, our cohort has shown M-F interface positivity in the absence of fetal tissue or AF positivity. This may highlight the importance of AF, or the fetal immune system, in protecting against fetal cCMV infection. Concurrent with this finding, one study containing 68 pregnant women with primary CMV infections reported false positive results from CMV PCR of AF. Within this group, 52 infants were CMV negative by urine culture, but 17 (33%) of the 52 had detectable CMV in the AF. The authors suggest that these results indicate virus transfer to the AF without fetal infection (41). The source of the early and transient detection of RhCMV DNA in the AF in four dams in our study is not clear (**Fig. 2A and 8A**). It is possible that RhCMV entry into the AF occurred during the viremic phase via intramembranous absorption, as has been described in rhesus macaques (33). Subsequently, the successful establishment of fetal infection may rely on several factors. Overall, the heterogenous spectrum of cCMV vertical transmission in humans and our model suggests CMV crosses a series of bottlenecks before accomplishing transmission or establishing fetal infection (**Fig. 8B**). These bottlenecks could be imposed by tissue barriers, as well as by circulating and tissue-resident maternal and fetal immune cell subsets.

We were also able to tease apart *in vivo* viral dynamics that highlighted the dominance of the clonal BAC-derived FL-RhCMV in the AF and fetal tissues of dam 001-101, despite apparently mixed populations in maternal plasma. Although it appears that FL-RhCMV was the dominant virus to cross the placenta and lead to widespread fetal infection in this dam, the clinical isolate RhCMV UCD52 did cross the placenta and lead to fetal infection in a previous study (16). We speculate that either FL-RhCMV has a competitive fitness advantage or, perhaps the placenta functions as a narrow bottleneck and FL-RhCMV established the initial infection in the fetal compartment. The latter contrasts with a previous estimate of bottleneck size at the M-F interface using only two genes, which suggested a wide bottleneck at the M-F interface (42). While comprehensive, this method for inferring sample-level strain frequencies is subject to several limitations. The approach used in this study used many SNV positions across the viral strain genomes to infer strain frequencies but was still limited by the need to select SNVs without strain bias and with sufficient sequencing coverage. Of note, our method for inferring strain frequency highlights variable nucleotide frequencies for each strain across this genome. Frequent recombination with multi-strain infections of HCMV is well-documented. We hypothesize that recombination between the two inoculating isolates may happen *in vivo*; however, assessing haplotypes and determining the extent to which recombination is occurring is outside the scope of this manuscript.

Sonographic measurements indicated that the presence of RhCMV at the M-F interface alone has an impact on fetal growth. Dams with higher PCR scores at the M-F interface had fetuses with smaller CRLs, compared to dams with lower PCR scores and compared to reference controls. Dams with high PCR scores also tended to have fetuses of lower weights at C-section. Surprisingly, fetal growth was not significantly impacted by AF DNA positivity or fetal tissue DNA positivity. In summary, abundance of RhCMV virus particles detected at the M-F interface negatively correlates with fetal weight and CRL in our cohort. This indicates that even when CMV is undetectable in AF or fetal tissues, the presence of CMV at the M-F interface can have an impact on fetal outcomes.

We were able to identify several candidate biomarkers of RhCMV vertical transmission. Fetal plasma TNF-alpha concentration was the strongest prognostic indicator differentiating AF+ and AF– dams. Our study also revealed lower levels of circulating BDNF in maternal plasma of AF+ dams as a unique biomarker previously not reported in the context of cCMV. Maternal BDNF is involved in placentation and placental health (43, 44), and clinical studies have shown that low maternal serum BDNF levels correlate with higher risk of low birth weight (45). Thus, sustained low levels of BDNF in AF+ dams may be predictive of placental or fetal pathology. Additionally, lower IL-10 in the maternal plasma and higher IL-21 and IFN-alpha in the AF at term gestation point to an increased inflammatory profile in the AF+ dams. This is corroborated by clinical studies demonstrating that CMV infection can lead to a bias of pro-inflammatory cytokines in the AF (46). These findings indicate the presence of increased pro-inflammatory and anti-viral responses within the fetuses exposed to CMV infection in utero.

Overall, this novel model of primary cCMV infection revealed a complex spectrum of vertical transmission, suggesting a larger population of newborns may be impacted by vertical transmission of CMV than previously appreciated. This highlights the need for multimodal detection when screening for cCMV to better define susceptible populations. Additionally, our analyses of maternal plasma, AF, and fetal plasma revealed potential biomarkers of CMV intrauterine transmission. The data generated herein highlight the translational significance of this model and demonstrates its potential for informing HCMV vaccine design and screening improvements.

Limitation of this study include the small number of animals. Larger studies will be needed to validate the predictive value of some of the biomarkers elucidated in this study for cCMV transmission. Another limitation is related to the sequencing analysis for inferring sample-level RhCMV strain frequencies and recombination events in maternal and fetal compartments. Future studies of viral sequences across dam and fetal anatomic compartments will enable in-depth studies of viral dynamics at the M-F interface. In future studies, this model can be leveraged to investigate immune cell subsets in the mother, M-F interface, and fetus that drive bottlenecks of CMV vertical transmission. Further, this model can test the efficacy of CMV vaccines to protect against fetal CMV transmission and disease, which is not possible in human clinical trials due to the overall low incidence, providing a critical pre-clinical assessment of prevention of symptomatic cCMV infection, a primary goal of a CMV vaccine.

## MATERIALS AND METHODS

### Subhead 1: Experimental Design

Animals in this study are RhCMV-seronegative, immunocompetent female rhesus macaques (*Macaca Mulatta*) breeding dams of Indian origin housed in the expanded specific pathogen-free (eSPF) colony at the Tulane National Primate Research Center (TNPRC, Covington, La). At the start of breeding season, colony-housed dams underwent periodic screening for pregnancy by abdominal ultrasound. Upon detection of pregnancy, 12 dams (aged 2.8-9.6 years) were enrolled in the study. All animal procedures were approved by the Institutional Animal Care and Use Committee (IACUC) at TNPRC. Animals were cared for in accordance with the NRC *Guide for the Care and Use of Laboratory Animals* and the Animal Welfare Act.

Between gestation days 55 to 69 (mean=61.6 days), enrolled dams were inoculated intravenously with two RhCMV strains. Pregnancies were monitored until approximately 21 weeks of gestation (11-12 weeks post-infection), at which point elective C-section was performed. Maternal blood was collected in EDTA vacutainer on days zero, one, four, seven, and then weekly until C-section. C-sections were performed at 19-22 weeks gestation (near term gestation). Saliva, urine, and AF were collected at assignment and thereafter weekly from day 0 pre-infection until C-section. Weekly ultrasounds were performed to monitor fetal biometrics.

At C-section, maternal blood, saliva, urine and AF were collected, fetus and placenta were removed from uterus, and decidua parietalis was swabbed from the uterus with surgical gauze. The cord was clamped on both sides and severed. Cord blood was collected from the umbilical vein into EDTA vacutainer from the placental facing cord to mimic procedure in humans. Fetal blood was collected by cardiac puncture. At fetal necropsy, fetal and placental morphometrics were recorded and all fetal tissues were collected for histopathological and virologic evaluation. Additionally, fetal urine was collected by cystocentesis.

### Subhead 2: Virus Inoculations

Dams were intravenously inoculated with a single dose of 1×10^6^ pfu RhCMV UCD52 and a single dose of 1×10^6^ pfu of Full length RhCMV (FL-RhCMV) diluted in one mL of sterile PBS and administered on opposite sides of the body. RhCMV UCD52 was chosen because it was previously found to be the dominant variant detected in plasma and AF following inoculation with a panel of three RhCMV variants (16). The FL-RhCMV strain is a BAC-derived full-length wild-type-like RhCMV clone (47). Challenge with both viruses mirrors a mix of RhCMV variants similar to natural infection with a virus swarm.

### Subhead 3: DNA Extraction and Measurement of RhCMV DNA by Real-time qPCR

DNA was extracted from maternal plasma, AF, maternal urine, fetal urine, maternal saliva, fetal tissues, and M-F interface tissues as described in supplemental methods. Absolute quantification of RhCMV DNA in tissues and fluids was determined by quantitative real-time PCR as previously described (21, 48) and detailed in supplementary methods. Maternal plasma, urine, and saliva samples, as well as maternal or fetal tissue samples were run in six technical replicates, while AF and fetal urine samples were run in 12 technical replicates. RhCMV DNA PCR on fetal or M-F interface tissue samples showing one of six replicates positive were repeated for another six technical replicates. At least two of 12 PCR replicates scoring positive at one or more copies in the PCR reaction were required to consider RhCMV DNA positivity in AF, M-F interface tissue and fetal tissue. RhCMV DNA in plasma and AF was reported as DNA copies per mL of sample, whereas PCR data on saliva, urine and tissues was reported as DNA copies per microgram of input DNA.

### Subhead 4: Placental RhCMV burden

Following real-time qPCR of individual components of the M-F interface tissues (**Fig. 2C**), we calculated one PCR score for the entire M-F interface of each dam as a metric of placental RhCMV burden. To generate this score, the mean number of positive PCR reads of all M-F interface tissues were added together and divided by the number of M-F interface tissues collected from that dam. We then calculated the average PCR score of the cohort (avg PCR score=4). By comparing individual PCR scores to the average PCR score of the cohort, six dams were categorized as PCR^lo^ (score <4), and six were categorized as PCR^hi^ (score >4). This metric was used as one of the grouping parameters to compare the effect of high vs low placental RhCMV burden on fetal crown-rump length and weight (**Fig. 4B-C**).

### Subhead 5: RNAScope *in situ* Hybridization

The assay was run using ACD RNAScope VS Universal Sample Prep and HRP Detection Reagents (Catalog# 323740 and 323210) using RhCMV and DapB probes (Catalog# 435299 and 320759). The RhCMV Probe targets the ORF for rh38, rh39, and rh55. Slides were digitally imaged at 40X with a Hamamatsu NanoZoomer360. CMV ISH positive cells were counted using a deep learning algorithm (HALO AI, Indica Labs) trained and then manually checked for accuracy by a pathologist. The deep learning algorithm was trained on training set of positive and negative control slides. Annotation regions were then drawn around entire tissue sections. The deep learning algorithm identified CMV positive events, and the pathologist visually confirmed or rejected based on location, size, and staining intensity. The number of CMV positive cells were reported over the total tissue are examined as CMV+ cells per cm2.

### Subhead 6: Fetal and placental measurements

Ultrasounds were performed weekly to monitor biparietal diameter and femur length using Edan Acclarix X8. Biometrics were recorded in duplicate and reported as mean mm. Gestational age was determined by sonographic measurements of the gestational sac diameter, as well as the crown-rump length. At C-section, ultrasound was performed to screen for characteristic cCMV abnormalities in brain and calcification in different organs, including brain and liver. These include periventricular calcifications, ventricular dilation, white matter gliosis, cerebral atrophy, cyst formations, abnormal sulcation, and cerebellum abnormalities. Fetal and placental weights were recorded postmortem. Fetal morphometric measures were performed with a caliper and measuring tape of the following metrics: abdominal circumference, crown rump length, head circumference, right foot, right hand, thoracic circumference, and transtemporal circumference.

### Subhead 7: Ultradeep Sequencing of RhCMV

DNA isolation was performed as described in the supplementary methods. Based on viral load, we used minimum 1000 input copies for amplification and library preparation, where available. AF at 82 days PI had approximately 300 input copies, while plasma at 77 and 82 days PI had fewer than 100 input copies. Sample DNA was non-specifically amplified using multiple displacement amplification (MDA, Repli-g, Qiagen #150025) then purified and concentrated using Zymo Genomic DNA Clean and Concentrate kit (Zymo Research, D4011). Sample DNA was equally split across 96 wells, each containing a single primer pair to span the entire genome and PCR amplified using the pure PfuUltra II Fusion HS DNA Polymerase (Agilent #600674). Primer sequences are listed in **Suppl Table 10**. Amplicons were pooled together then purified using QIAprep spin mini prep kit (Qiagen #27106). Using 500 ng of purified DNA, library prep was performed using NEB FS DNA library prep kit (E7805) and dual-index Illumina adapters. Libraries were run on Illumina Nextseq 2000 platform using a 200-cycle cartridge.

De-multiplexed reads with adapter sequences removed were loaded into CLC Genomics workbench and processed using a modified version of Qiagen’s SARS-CoV-2 ARTIC V3 analysis pipeline. Reads were trimmed by base quality and separately mapped to either FL-RhCMV (GCA_027929765.1) or RhCMV UCD52 (GCA_027930255.1) reference genomes. Due to limited consensus-level differences (two SNVs or indels in each reference) in the inoculation stock compared to the respective reference genomes, the published reference genomes were used as reference sequences in downstream analyses. Read mapping was refined by removing ligation artifacts, performing local realignment based on structural variant calls, removing reads with unaligned ends, and trimming bases aligning to the primer sequences at the ends of reads. Output BAM files for each sample were used in downstream analyses.

To infer strain frequencies, read depths were calculated for all samples using samtools v1.19.2 (49) function mpileup with additional flags to filter low quality bases and allow for character parsing: --min-BQ 20, --max-depth 0, --no-output-ends, --no-output-ins, and --no-output-del. Next, each reference genome was aligned to the other, and differing positions were extracted using NucDiff v2.0.2 (50). Outputs (ref_snps.gff) were subset to only include SNVs. Insertions, deletions, and multiple nucleotide variants (MNVs) were not considered. Filter schema is shown in **Fig. S5** and total number of remaining SNVs used for each sample-level strain frequency calculation can be found in **Suppl Table 4**. Finally, strain frequencies at each remaining SNV position for each sample were calculated by dividing the number of reads with a nucleotide matching the reference genome (i.e. RhCMV UCD52 or FL RhCMV) by total read depth at that position. Mean frequencies across all positions were also calculated for each sample. We performed the above procedure using both reference genomes.

### Subhead 8: Luminex

Luminex assay was used to determine cytokine levels in the maternal plasma, fetal plasma, cord blood, and AF. We used the R&D Systems NHP XL Cytokine Premixed 33-plex Panel (Cat# FCSTM21-33). EDTA plasma and AF samples were analyzed in two replicates and were prepared using a 1:2 dilution. To optimize sample dilution needed for linearity, serial dilutions was performed and 1:2 dilution in PBS was determined. Reactions were read using the Bio-Plex® 200 System (Bio-Rad Laboratories, Hercules, CA), and results were calculated on the Bio-Plex Manager™ Software (Bio-Rad) at the TNPRC Pathogen Detection and Quantification Core. Mean concentrations of replicates were plotted. We included values which were estimated based on extrapolation from the standard curve. We excluded values from wells that had a low bead count (below 50 beads) or a high %CV (above 30%).

### Subhead 9: Statistical Methods

Functional mean trajectory of each CBC panel component was smoothed via a Gaussian kernel using fdapace package in R (51). Durbin test within coin package was used to test for association of time with CBC, and separately with Luminex maternal plasma data during acute phase. Statistical significance was determined via Monte-Carlo (10000 permutations). Multiple testing correction was done using FDR (52) with significance set at an adjusted p-value less than 0.2. For markers and analytes that demonstrated significant difference by time, we performed three and four follow up tests, respectively. CBC markers compared day seven, 21, and 84 to day zero, while analytes compared day two, four, seven, and 14 back to day zero via exact Wilcoxon signed rank test using coin package in R (53). Due to inconsistent timing of CBC measures at week 12, values recorded between day 81–84 were used for change from baseline at day 84. In these post-hoc tests, multiple testing correction was done via Holm procedure.

Global differences in fetal morphometric parameter z-scores between groups were assessed via non-parametric MANOVA (multivariate analysis of variance) using npmv package in R (54). Fetal morphometric parameter z-scores were derived as gestational age-specific mean-centered and standard deviation-scaled values using published population estimates (23). Missing z-scores were imputed with animal specific median z-score of observed z-score parameters. Exact Wilcoxon rank-sum tests were conducted to determine differences between groups in each morphometric parameter and for comparing PE in our cohort to gestation-comparable controls (53). Binomial tests on proportions determined whether median z-scores and proportion below small for gestational age (10th percentile) of the population of macaques under study differed from the referent population for crown-rump length and fetal weight.

AUC was calculated using the standard trapezoid rule for analyzing viral loads in maternal plasma, urine, and saliva calculating cumulative viral load over different time frames. If an animal lacked a day zero observation, they were dropped from analysis. If they lacked a day 14 observation, we carried forward the last observation. We performed a similar procedure for calculating AUC over the acute plasma Luminex data (first 14 days). We tested for differences in these AUC values as well as other Luminex data (C-section, fetal plasma, etc) by AF status via exact Wilcoxon rank sum test. Multiple testing correction was done separately within each sample source/analysis (i.e. VL and acute plasma were separate corrections) with significance set at adjusted p-value less than 0.2. We set data below the limit of detection to the lower limit divided by 2.

We performed PCA separately on data from fetal plasma, AUC of the maternal acute plasma, C-section, cord blood, and maternal plasma. For PCA visualization, missing values were replaced with the mean value for that analyte. Values below limit of detection were again set to the lower limit divided by 2. We scaled and centered the data prior to calculating principal components. Survival analyses comparing either placental transmission or fetal survival between immunocompetent and CD4 depleted animals was done via log-rank test using R package survival. We tested whether Spearman correlation between ISH and PCR data within animal 001-101. Exact Wilcoxon rank sum tests were performed for placental efficiency and chemerin analyses.

## Supplementary Materials

**Materials and Methods**

**Supplementary Figures**

Fig. S1 to S8.

**Supplementary Tables**

Table S1 to S10.

**Data File**

All data.xlx

**References** (55–57)

## Acknowledgements

The authors acknowledge the Veterinary Medicine staff at the Tulane National Primate Research Center (TNPRC) for their expert care of the animals, and the following TNPRC Core services: Clinical Pathology Core (RRID:SCR_024609) for complete blood count analysis; Anatomic Pathology Core (RRID:SCR_024606) and Confocal Microscopy and Molecular Pathology Core (RRID:SCR_024613) for histopathology, image analysis and RNAScope services; and the Pathogen Detection and Quantification Core for performing and reading Luminex assays (RRID:SCR_024614).

## Funding

National Institutes of Health grant P01 AI129859

TNPRC Base Grant P51OD011104 (RRID: SCR_008167)

S10 OD026800

## Author Contributions

Conceptualization: TM, MM, DM, KF, SRP, RVB, AK.

Methodology: TM, MM, CMC, RVB, AK.

Validation: TM, MM, ES, CMC, ML, LR, AM, TK, RB, AD.

Formal analysis and interpretation: TM, MM, ES, RVB, RB, AD, CC, ML, LR, JG, SL, CEO.

Investigation: TM, MM, ES, SK, MT, FB, LS, CEO, CCM, AM, RVB.

Data Curation: TM, MM, SK, ES, RVB, RB, AD, CMC, AM.

Writing- original draft preparation: TM, RB, AK.

Writing- review and editing: All authors.

Visualization: TM, MM, CMC, CEO, AD, RB, RVB.

Resources: MS, VD, LDM, DM, KF, RB, AD, CC, CMC, ML, LR, JG, KO, SL.

Supervision: KF, TK, CC, SRP, AK.

Project Administration: TM, MM, ES, AK.

Funding Acquisition: KF, SRP, AK.

## Competing Interests

### Data and Materials Availability

https://github.com/cmc0043/rhcmv-inferred-strain-frequency

## Notes

https://github.com/cmc0043/rhcmv-inferred-strain-frequency

### Competing Interest Statement

The authors have declared no competing interest.

