## Supplementary Methods for "A nonhuman primate model mirrors human congenital cytomegalovirus infection and reveals a spectrum of vertical transmission outcomes"

**Supplemental Methods**

**Subhead 1: Sample Collection and Processing**

Saliva, urine, AF, and PBMCs were collected weekly from each dam enrolled in the study as previously described (*16*) (*18*). Saliva was collected by oral saline wash, concentrated using Ultracel YM-30 (Amicon/Milipore), then aliquoted for storage at -20C. Urine samples were collected by clean pan catch, cellular material and debris was pelleted, and the supernatant was concentrated using Ultracel YM-30 (Amicon/Milipore), then aliquoted for storage at -20C. AF was collected by amniocentesis, centrifuged to remove debris, and supernatant was aliquoted for storage at -20C. Following plasma collection, PBMCs were isolated by ficoll separation using Lymphocyte Separation Media (LSM) (MP Biomedicals). All PBMCs were cryopreserved using serum-free freezing media Bambanker BB01 (Bulldog-Bio).

While working under sterile practice, placental tissues were harvested at C-section into cold PBS and at fetal necropsy into cold R10 media. The placenta was first processed in necropsy for top-down pictures of maternal and fetal side. In addition, sections of full-thickness placenta, cord and amniotic membrane were placed zinc-formalin fixative and paraffin-embedded for preservation for H&E and in situ hybridization. Remaining placenta was cold transported to research lab for further processing. The primary and, if present, the secondary placenta were washed in a petri dish which was place over wet ice with saline solution to remove maternal blood, and visible blood clots were removed. After extensive cleaning (~1 hour), tissues were dissected as follows for snap freezing with liquid nitrogen. The extensive cleaning is important for both viability and to ensure tissue-bound cells are not investigated. Cord was cut in 1cm pieces, amniotic membrane was cut in 1-2cm^2 squares from a location far from the placenta discs. The decidua layer of placenta was removed using forceps to pull of the tissue layer of decidua basalis from the maternal side of the placenta(s). Care was taken to not pull off the epithelial layer of the placenta (too deep). Sterile gauze used to wipe the uterus of both decidua parietalis and basalis from surgery was moved one by one into the lid of the petri dish and screened for decidua tissue which was moved to a petri dish with R10 media on wet ice. After cleaning and decidua collection, each placental disc was cut with a scalpel into a 2x2 cm^2 away from the margin and moved to a separate round petri dish with 1X PBS. This full thickness (excluding decidua) of placenta was cut into 0.5cm wide strips and flipped on its side. The maternal side and fetal side were dissected apart and placed in a 1.5mL snap cap tube for snap freezing of maternal and fetal side. This was repeated for both the primary and if present also for the secondary placenta. The zone between maternal and fetal size was excluded to not risk cross-contamination. The decidua was further cleaned for any small remaining blood clots in a petri dish and media was replaced with 1X PBS to easily spot any tiny blood clots. Clean decidua pieces of about 0.5-1cm in size were snap frozen. The remaining tissue was washed in 50mL conical with ~40mL of saline into a course metal strainer. Washed tissue was transferred with a backward rinse from the metal strainer via a funnel back into a clean 50mL conical. This process was repeated 4X to prepare washed decidua tissue. Washed decidua was transferred back to a petri dish and resuspended with 1X PBS to cover the tissue for further dissection. Using a scalpel 3-4mm pieces of decidua was prepared and then transferred into a 25cm$^2$ cell culture flask for enzymatic digestion.

Complete fetal necropsies were also performed and included sampling of 42 tissues per fetus. Collection focused on tissues with known CMV-tropism (brain, lung, kidney, cochlea). processed for lymphocyte isolation, snap frozen for DNA extraction, and placed in zinc-formalin and processed routinely for paraffin blocks. Tissues sections were cut into 4 um sections and mounted on charged slides prior to staining (H&E or in situ hybridization). DNA was extracted from 1-25 mg of snap-frozen tissue using DNeasy Blood and Tissue kit (Qiagen).

**Subhead 2: Complete Blood Count**

For each dam, whole blood was collected on days zero, one, seven, and weekly PI until time of C-section. A pre timepoint was also collected when possible, considering colony logistics, housing acclimation period, and timing of pregnancy detected. The TNPRC clinical lab assessed the whole blood samples using a Sysmex XXN-1000v analyzer, which allows for determination of complete blood counts (CBC). This provides absolute counts of granulocytes, lymphocytes, and monocytes. Abnormal counts are reviewed by a manual differential.

**Subhead 3: DNA Extraction and Measurement of RhCMV DNA by Real-time qPCR**

Qiagen Kit QIAamp DNA Blood Mini Kit (catalog# 55106) was used to extract DNA from plasma, saliva, and AF. Qiagen Kit QIAamp Viral RNA Mini Kit (catalog# 52906) was used to extract DNA from 140 uL of urine. DNA was extracted from 200 uL of plasma, then each plasma sample was eluted in 150 uL of elution buffer. DNA from AF, saliva, or urine was extracted and eluted in 100 uL of elution buffer. DNA was extracted from maternal and fetal tissues using Qiagen QIAamp Fast DNA Tissue Kit (catalog# 51404). Snap-frozen tissues were kept chilled with dry ice and dissected in a petri dish. DNA was extracted from 1-25mg of snap-frozen tissue using DNeasy Blood and Tissue kit (Qiagen).

Absolute quantification of RhCMV DNA in tissues and fluids was determined by quantitative real-time PCR as previously described (*48*) (*21*). The Forward Primer 5'-GTTTAGGGAACCGCCATTCTG-3', Reverse primer 5'-GTATCCGCGTTCCAATGCA-3', and Probe 5'-FAM-TCCAGCCTCCATAGCCGGGAAGG-TAMRA-3' added to a 25 μL reaction with Supermix Platinum Quantitative PCR SuperMix-UDG (Invitrogen). Primers and probes target conserved noncoding exon 1 region of the immediate early gene and have been used to amplify different RhCMV strains (*47*). Reaction was performed on a 96-well plate using QuantStudio6 (ThermoFisher Scientific). To control for non-target sequence lowering efficiency of reaction, a screened cohort of RhCMV-seronegative animals PBMC's were DNA extracted and used as seronegative rhesus genomic DNA. This DNA is screened by the same assay to confirm genomic DNA as negative. Standard curve target sequence is diluted in 3 ng/uL seronegative genomic DNA. The lower limit of detection of this assay is between one to ten RhCMV DNA copies in the PCR reaction.

**Subhead 4: RNAscope In Situ Hybridization Tissue Pretreatment**

Four $u$m tissue sections were mounted on Superfrost Plus Microscope slides, baked for two hours at 60$^o$C and passed through three changes of Xylene and three changes of 100% ethanol to remove paraffin. Slides were air dried, labeled, and loaded onto the Ventana Discovery Ultra autostainer. The assay was run using ACD RNAScope VS Universal Sample Prep and HRP Detection Reagents (Catalog# 323740 and 323210). Tissue pretreatments include Target Retrieval at 97$^o$C for 16 or 24 minutes, depending on the tissue, and protease at 37$^o$C for 16 minutes. The RhCMV and DapB probes (Catalog# 435299 and 320759) were incubated for two hours at 43$^o$C. Green color development was done at the recommended time of 20 minutes, followed by counterstaining with hematoxylin II and bluing reagent at 16 minutes each. Upon completion, slides were removed and put through alternating manual washes of deionized water containing 0.1% Dawn dish soap and plain deionized water for a total of five cycles. Slides were then cleared in ethanol (80%, 95%, 100%, 100%) and three xylene changes before being permanently mounted with StatLab Acrymount mounting media. Slides were then dried overnight.

**Subhead 5: Ultradeep Sequencing of RhCMV**

Hypervariable regions were identified by calculating the number of SNVs within 100 bp windows across each reference genome. SNVs within regions that had >5 mean differences within 100 bps were removed to reduce the impact of reference genome mapping bias. SNVs with low coverage (<250 reads) in the non-reference, sequenced inoculum sample (e.g. sequenced FL RhCMV inoculum mapped to RhCMV UCD52 reference genome) were also removed. To account for heterogeneity in the stocks, particularly in the passaged RhCMV UCD52 isolate, positions in each stock that had >1% frequency of the alternate stock were filtered out. In total, the filters removed 2015 of 2865 SNVs identified as differing between the two reference genomes with RhCMV UCD52 as the reference and 2000 of 2862 SNVs with FL-RhCMV as the reference. At the sample level, SNVs with coverage of <100 reads were also removed to avoid estimating strain frequencies in genomic positions with few aligned reads.

Custom scripts were generated in R Statistical Environment (*55*) with extension packages from Comprehensive R Archive Network (CRAN; https://cran.r-project.org/), including tidyverse v2.0.0 (*56*), and knitr v1.45 (*57*) to summarize the allele depth corresponding to each strain at each position, apply reference strain SNV filters to account for mapping bias, and calculate inferred strain frequencies based on the mean frequency of alleles corresponding to each reference strain at each resulting SNV position. The code for replicating these analyses is available through a public source code repository (https://github.com/cmc0043/rhcmv-inferred-strain-frequency /).

**Subhead 6: IgM Serology of Fetal Plasma**

Fetal IgM responses were assessed using fetal plasma collected at C-section. Fetal plasma was chosen over cord blood due to potential contribution of maternal blood in cord blood samples. Plasma was diluted 1:5 with sterile PBS (25 µL plasma plus 225 µL PBS) and incubated overnight in a Protein A trap plate (Cytiva Cat# 28903133) supplemented with 100 µL of Protein G agarose resin (Pierce) at 4°C with constant agitation to remove IgG from the plasma to reduce potential background and competition for antigen binding with IgM in our assay. The trap plate was spun at 200 x g for one minute with a 96 well round bottom plate (Corning) attached to collect the flow-through containing IgG-depleted plasma, and each well was washed once more with an equivalent volume (250 µL) of PBS. IgG-depletion was confirmed using a total IgG sandwich ELISA, using unlabeled and HRP-conjugated mouse anti-monkey IgG (Clone SB108a, Southern Biotech) for coating and detection, respectively. For detection of RhCMV-specific IgM in IgG-depleted plasma, we performed a whole virion ELISA as previously described (*17*). Briefly, high binding 384-well clear plates (Corning) were coated overnight at 4°C with 2000 $TCID_{50}$/mL UCD52 or 8000 $TCID_{50}$/mL FL- RhCMV diluted in 0.1M sodium bicarbonate buffer (pH=9.55). After blocking for one to two hours at ambient temperature, we added serially diluted plasma samples and incubated for one to two hours at ambient temperature. IgM binding was detected using a goat anti-monkey IgM mu chain-HRP antibody (Rockland cat. 617-101-007) incubated for one hour at 1:8000. TMB substrate (TMB SureBlue, KPL) was used to develop plates for seven minutes, followed by 0.1% HCl (TMB Stop, KPL) to stop the reaction, and plates were read at 450 nm on a spectrophotometer (BioTek Synergy). Results are reported as the OD450 of undiluted IgG-depleted plasma (1:10 from original plasma sample), and a positivity cutoff was established at 2 standard deviations above the mean $OD_{450}$ of two control animals with RhCMV-seronegative plasma samples.

**Subhead 7: Chemerin Immunoassay**

Amniotic fluid chemerin levels were measured using the quantitative sandwich enzyme immunoassay (Quantikine ELISA) by R&D Systems (Catalog# DCHM00). Before conducting the assay, we used NCBI Blast to confirm that human and Rhesus macaque chemerin (NCBI reference genome Mmul_10) have 100% homology to ensure the assay is appropriate for our model. The assay was conducted according to the manufacturer's instructions. All samples were run in duplicate and read on a Synergy 2 (BioTek) microplate reader at 450nm wavelength.
