## Supplementary Tables for "A nonhuman primate model mirrors human congenital cytomegalovirus infection and reveals a spectrum of vertical transmission outcomes"

**Supplementary Table 1. Characteristics of RhCMV-Seronegative Female Macaques Enrolled in Study**

| Animal ID | Dam Age (Yrs) | Number of Prior Pregnancies | GD at RhCMV Infection | GD at C-section | Weeks PI at C-section | Sex of Fetus |
|---|---|---|---|---|---|---|
| **001-101** | 9.6 | 3 | 67 | 149 | 11 | F |
| **001-102** | 8.7 | 1 | 69 | 154 | 12 | F |
| **001-103** | 6.6 | 1 | 56 | 139 | 11 | F |
| **001-104** | 7.4 | 3 | 67 | 138 | 11 | F |
| **001-105** | 8.5 | 3 | 67 | 149 | 11 | M |
| **001-106** | 5.7 | 1 | 60 | 142 | 11 | F |
| **001-107** | 6.5 | 2 | 65 | 153 | 12 | F |
| **001-108** | 6.4 | 1 | 57 | 146 | 12 | M |
| **001-109** | 5.6 | 1 | 60 | 143 | 11 | F |
| **001-110** | 3.9 | 1 | 57 | 140 | 11 | F |
| **001-111** | 3.9 | 0 | 59 | 142 | 11 | F |
| **001-112** | 2.8 | 0 | 55 | 136 | 11 | M |

*Gestation day (GD)

*Post-infection (PI)

**Supplementary Table 2. Placental lesions found in dams at the time of C-section.**

| Animal ID | Placentitis | Chorioamnionitis | Placental Infarct |
|---|---|---|---|
| **001-102** | X | | |
| **001-104** | | X | |
| **001-106** | X | | |
| **001-110** | X | | |
| **001-111** | X | | X |
| **001-112** | | | X |

**Supplemental Table 3. Fetal Morphometric Comparisons in RhCMV-Infected Dams.**

**A)** **Median (IQR)**

| Groups | Strata | n | Abdominal circumference (cm) | Crown-rump length (cm) | Fetal weight (g) | Head circumference (cm) | Right foot (cm) | Right hand (cm) | Thoracic circum-ference (cm) | Total Placenta weight (g) | Trans-temporal (cm) |
|---|---|---|---|---|---|---|---|---|---|---|---|
| Amniotic Fluid | PCR Negative | 7 | -0.7 (-0.7, 0.2) | -0.8 (-1.7, 0.0) | -0.2 (-0.7, 0.2) | -0.5 (-0.9, 0.1) | -0.6 (-0.9, 0.1) | -1.3 (-1.9, 0.0) | -0.6 (-1.7, 0.3) | 0.8 (0.4, 1.3) | -0.5 (-0.9, 0.7) |
| | PCR Positive | 5 | -0.3 (-0.4, 0.0) | -1.7 (-3.2, -0.6) | -0.8 (-1.3, -0.1) | 0.1 (-0.2, 0.2) | -0.4 (-0.4, -0.3) | -0.9 (-1.3, -0.6) | -0.1 (-0.3, 0.1) | 0.6 (0.1, 0.9) | -0.8 (-1.5, -0.6) |
| Fetal Tissue | PCR Negative | 9 | -0.5 (-0.7, 0.6) | -0.8 (-2.8, 0.7) | -0.1 (-0.7, 0.4) | -0.3 (-0.6, 0.1) | -0.3 (-1.0, 0.4) | -1.3 (-1.7, 0.0) | -0.1 (-1.1, 0.3) | 0.8 (0.1, 1.2) | -0.6 (-1.5, 0.4) |
| | PCR Positive | 3 | -0.2 (-0.4, -0.2) | -1.4 (-1.6, -1.0) | -1.0 (-1.1, -0.7) | 0.2 (-0.3, 0.4) | -0.4 (-0.5, -0.4) | -0.9 (-1.5, -0.7) | -0.3 (-0.5, 0.1) | 0.6 (0.3, 0.9) | -0.8 (-0.9, -0.3) |
| Maternal-Fetal Interface | PCR Lo | 6 | -0.4 (-0.7, 0.5) | -0.6 (-1.2, 0.3) | -0.3 (-0.4, 0.2) | 0.1 (-0.7, 0.3) | -0.4 (-0.5, 0.2) | -0.3 (-1.1, 0.0) | 0.1 (-0.5, 0.4) | 1.0 (0.3, 1.2) | -0.7 (-0.8, 0.2) |
| | PCR Hi | 6 | -0.4 (-0.7, -0.2) | -2.3 (-3.1, -1.0) | -1.2 (-1.3, -0.1) | -0.3 (-0.6, -0.2) | -0.7 (-1.1, -0.3) | -1.5 (-2.1, -1.0) | -0.3 (-0.6, -0.3) | 0.7 (-0.1, 0.8) | -1.0 (-1.5, 0.0) |

**B)**

| Groupings | Abdominal circumference (cm) | Crown-rump length (cm) | Fetal weight (g) | Head circumference (cm) | Right foot (cm) | Right hand (cm) | Thoracic circumference (cm) | Total Placenta weight (g) | Transtemporal (cm) |
|---|---|---|---|---|---|---|---|---|---|
| AF PCR | 0.807 | 0.370 | 0.570 | 0.167 | 0.684 | 0.935 | 0.744 | 0.416 | 0.328 |
| Fetus PCR | 0.517 | >0.999 | 0.229 | 0.309 | 0.643 | 0.711 | 0.781 | 0.643 | 0.853 |
| Placenta PCR | 0.297 | 0.092 | 0.337 | 0.337 | 0.378 | 0.297 | 0.377 | 0.336 | 0.422 |

### Supplemental Table 4. Single Nucleotide Polymorphisms (SNPs) in Sequenced Samples.

| Sample | Mapping reference | SNPs used for analysis | UCD52 mean strain frequency (SE) | FL mean strain frequency (SE) | Other mean strain frequency (SE) |
|---|---|---|---|---|---|
| 965BACmda | FL-RhCMV | 862 | 0.09321 (0.005581) | 99.88 (0.005627) | 0.02423 (0.001002) |
| 965BACmda | UCD52 | 850 | 0.1115 (0.005846) | 99.86 (0.005992) | 0.02685 (0.001453) |
| Dam 001-101 AF77 | FL-RhCMV | 775 | 0.02026 (0.002829) | 99.96 (0.0038) | 0.02013 (0.002621) |
| Dam 001-101 AF77 | UCD52 | 764 | 0.04041 (0.005267) | 99.94 (0.00624) | 0.0221 (0.003469) |
| Dam 001-101 AF82 | FL-RhCMV | 575 | 0.0214 (0.002817) | 99.96 (0.004068) | 0.02204 (0.002369) |
| Dam 001-101 AF82 | UCD52 | 565 | 0.03156 (0.003407) | 99.94 (0.004217) | 0.0235 (0.002651) |
| Dam 001-101 fetal basal ganglion | FL-RhCMV | 602 | 0.01563 (0.002518) | 99.97 (0.00263) | 0.0113 (0.0008016) |
| Dam 001-101 fetal basal ganglion | UCD52 | 591 | 0.03065 (0.005965) | 99.96 (0.00599) | 0.01153 (0.00083) |
| Dam 001-101 fetal heart | FL-RhCMV | 669 | 0.03007 (0.003826) | 99.95 (0.004236) | 0.01645 (0.001626) |
| Dam 001-101 fetal heart | UCD52 | 660 | 0.04444 (0.004658) | 99.94 (0.005171) | 0.01807 (0.002142) |
| Dam 001-101 fetal kidney | FL-RhCMV | 664 | 0.01373 (0.001487) | 99.97 (0.001823) | 0.01431 (0.001148) |
| Dam 001-101 fetal kidney | UCD52 | 656 | 0.02765 (0.003205) | 99.96 (0.003746) | 0.01683 (0.001978) |
| Dam 001-101 fetal lung | FL-RhCMV | 661 | 0.01924 (0.002428) | 99.96 (0.006714) | 0.02522 (0.00633) |
| Dam 001-101 fetal lung | UCD52 | 647 | 0.03987 (0.005094) | 99.93 (0.008116) | 0.02705 (0.006492) |
| Dam 001-101 P14 | FL-RhCMV | 793 | 60.17 (0.857) | 39.81 (0.8571) | 0.02391 (0.001184) |
| Dam 001-101 P14 | UCD52 | 780 | 61.23 (0.8503) | 38.74 (0.8504) | 0.02409 (0.001479) |
| Dam 001-101 P4 | FL-RhCMV | 794 | 9.272 (0.637) | 90.71 (0.637) | 0.01991 (0.001724) |
| Dam 001-101 P4 | UCD52 | 782 | 9.612 (0.6555) | 90.37 (0.6555) | 0.01946 (0.001292) |
| Dam 001-101 P7 | FL-RhCMV | 760 | 29.33 (1.201) | 70.65 (1.201) | 0.01967 (0.001358) |
| Dam 001-101 P7 | UCD52 | 751 | 30.34 (1.221) | 69.64 (1.221) | 0.01839 (0.0009919) |
| Dam 001-101 P77 | FL-RhCMV | 14 | 0.03547 (0.03547) | 99.94 (0.04298) | 0.02716 (0.02716) |
| Dam 001-101 P77 | UCD52 | 14 | 0.07392 (0.03901) | 99.87 (0.05566) | 0.05794 (0.03945) |
| Dam 001-101 P82 | FL-RhCMV | 273 | 59.08 (2.857) | 40.9 (2.856) | 0.01419 (0.001392) |
| Dam 001-101 P82 | UCD52 | 275 | 59.8 (2.84) | 40.19 (2.84) | 0.01386 (0.001381) |
| Dam 001-101 fetal parietal cortex | FL-RhCMV | 779 | 0.08491 (0.02818) | 99.9 (0.02823) | 0.01442 (0.001013) |
| Dam 001-101 fetal parietal cortex | UCD52 | 767 | 0.1034 (0.03004) | 99.88 (0.03009) | 0.01638 (0.001385) |
| UCD52mda | FL-RhCMV | 862 | 99.93 (0.003923) | 0.04987 (0.003876) | 0.01896 (0.0007992) |
| UCD52mda | UCD52 | 850 | 99.94 (0.003651) | 0.04014 (0.003608) | 0.01782 (0.0006893) |

**Supplemental Table 5. Change from baseline for individual analytes.**

| Analyte | P value | p.adj | Day 1 | Day 4 | Day 7 | Day 14 |
|---|---|---|---|---|---|---|
| IL-15 | <1e-4 | <1e-4 | **0.00781** | **0.00391** | **0.00781** | **0.00391** |
| PD-L1 | <1e-4 | <1e-4 | **0.0156** | **0.00586** | **0.0312** | **0.00391** |
| IP-10 | <1e-4 | <1e-4 | **0.0391** | **0.00293** | **0.0156** | **0.00195** |
| VEGF | <1e-4 | <1e-4 | 0.359 | **0.00781** | **0.0234** | 0.203 |
| MIP-3 alpha | 0.0042 | 0.007412 | 0.492 | **0.0483** | 0.297 | **0.0371** |
| PDGF-AA | <1e-4 | <1e-4 | 0.496 | **0.00293** | 0.0781 | **0.00195** |
| IL-6 | <1e-4 | <1e-4 | 0.875 | **0.00781** | **0.0234** | 0.875 |
| IFN-alpha | 0.0007 | 0.0015 | 1 | **0.0156** | 1 | 0.938 |
| IFN-gamma | <1e-4 | <1e-4 | 1 | **0.0156** | 0.375 | 0.5 |
| I-TAC | <1e-4 | <1e-4 | 0.75 | **0.0156** | 0.0938 | 0.75 |
| IL-10 | <1e-4 | <1e-4 | 0.75 | **0.00781** | 0.25 | 0.0938 |
| GM-CSF | <1e-4 | <1e-4 | 0.875 | **0.00391** | 0.188 | 0.406 |
| Granzyme B | 0.0232 | 0.03867 | 0.945 | 0.117 | 0.875 | **0.0156** |
| CXCL13 / BLC | 0.0003 | 0.00075 | 1 | 1 | 1 | 0.25 |
| CXCL2 / GRO beta | 0.0004 | 0.0009231 | 1 | 0.0625 | 0.656 | 0.656 |
| Eotaxin | 0.0001 | 0.0002727 | 1 | 0.168 | 0.938 | 1 |
| IFN-beta | 0.0339 | 0.05085 | 1 | 0.25 | 1 | 1 |
| CD40 Ligand | 0.003 | 0.005625 | 0.695 | 0.109 | 0.388 | 0.388 |
| BDNF | 0.0337 | 0.05085 | 0.835 | 0.835 | 0.438 | 0.835 |
| TNF-alpha | 0.0016 | 0.0032 | 0.875 | 0.0938 | 0.375 | 0.906 |
| G-CSF | 0.866 | 0.8959 | | | | |
| IL-12 p70 | 0.1838 | 0.2297 | | | | |
| IL-13 | 1 | 1 | | | | |
| IL-17 / IL-17A | 0.4226 | 0.4876 | | | | |
| IL-2 | 0.0832 | 0.1135 | | | | |
| IL-21 | 0.4752 | 0.528 | | | | |
| IL-4 | 0.3782 | 0.4538 | | | | |
| IL-7 | 0.0605 | 0.08643 | | | | |
| IL-8 | 0.6858 | 0.7348 | | | | |

| Analyte | P value | p.adj | Day 1 | Day 4 | Day 7 | Day 14 |
| --- | --- | --- | --- | --- | --- | --- |
| PDGF-BB | 0.1414 | 0.1844 | | | | |

**Supplemental Table 6. Luminex assay on maternal plasma at C-section.**

| Analyte | AF– Dams | AF+ Dams | P value | FDR |
|---|---|---|---|---|
| IL-10 | 39.5 (n=7) | 6.48 (n=5) | 0.0455 | 0.502 |
| BDNF | 2.88e+03 (n=7) | 1.61e+03 (n=5) | 0.048 | 0.502 |
| IP-10 | 198 (n=6) | 123 (n=5) | 0.0519 | 0.502 |
| IL-8 | 2.04e+03 (n=7) | 1.65e+03 (n=5) | 0.106 | 0.604 |
| IL-7 | 6.26 (n=6) | 1.41 (n=5) | 0.119 | 0.604 |
| TNF-alpha | 0.13 (n=7) | 2.66 (n=5) | 0.125 | 0.604 |
| IL-17 / IL-17A | 0.395 (n=7) | 0.395 (n=5) | 0.311 | 1.00 |
| G-CSF | 3.52 (n=7) | 3.52 (n=4) | 0.321 | 1.00 |
| Eotaxin | 327 (n=7) | 190 (n=5) | 0.323 | 1.00 |
| IL-6 | 0.875 (n=7) | 6.39 (n=5) | 0.413 | 1.00 |
| IFN-gamma | 2.15 (n=7) | 2.15 (n=5) | 0.417 | 1.00 |
| VEGF | 30.8 (n=7) | 25.9 (n=5) | 0.432 | 1.00 |
| IL-2 | 6.41 (n=7) | 6.36 (n=5) | 0.61 | 1.00 |
| IFN-beta | 3.25 (n=7) | 3.25 (n=5) | 0.691 | 1.00 |
| CXCL2 / GRO beta | 1.26e+03 (n=6) | 917 (n=4) | 0.762 | 1.00 |
| CD40 Ligand | 1.89e+03 (n=6) | 1.38e+03 (n=5) | 0.792 | 1.00 |
| PDGF-BB | 1.54e+03 (n=7) | 902 (n=3) | 0.833 | 1.00 |
| MIP-3 alpha | 35.7 (n=7) | 35.7 (n=5) | 0.876 | 1.00 |
| PDGF-AA | 55.3 (n=7) | 55.7 (n=5) | 0.876 | 1.00 |
| IFN-alpha | 0.01 (n=5) | 0.525 (n=4) | 0.905 | 1.00 |
| IL-15 | 3.76 (n=7) | 3.93 (n=5) | 0.912 | 1.00 |
| PD-L1 | 419 (n=6) | 477 (n=5) | 0.931 | 1.00 |
| GM-CSF | 8.22 (n=7) | 8.22 (n=5) | 0.965 | 1.00 |
| I-TAC | 1.18 (n=7) | 1.18 (n=5) | 1.00 | 1.00 |
| Granzyme B | 12.8 (n=7) | 11.1 (n=5) | 1.00 | 1.00 |
| IL-4 | 0 (n=7) | 0 (n=5) | 1.00 | 1.00 |
| IL-12 p70 | 0.7 (n=7) | 0.7 (n=5) | 1.00 | 1.00 |
| IL-21 | 0.37 (n=7) | 0.37 (n=5) | 1.00 | 1.00 |
| CXCL13 / BLC | 11.9 (n=7) | 11.9 (n=5) | 1.00 | 1.00 |

**Supplemental Table 7. Luminex assay on amniotic fluid at C-section.**

| Analyte | AF– Dams | AF+ Dams | P value | FDR |
|---|---|---|---|---|
| IFN-alpha | 5.62 (n=7) | 13.3 (n=5) | 0.0265 | 0.342 |
| IL-21 | 88.4 (n=7) | 132 (n=5) | 0.0467 | 0.342 |
| BDNF | 49.9 (n=7) | 37.8 (n=5) | 0.0732 | 0.342 |
| IFN-beta | 3.64e+03 (n=7) | 5.23e+03 (n=5) | 0.0732 | 0.342 |
| IFN-gamma | 357 (n=7) | 696 (n=5) | 0.0732 | 0.342 |
| VEGF | 660 (n=7) | 919 (n=5) | 0.0732 | 0.342 |
| MIP-3 alpha | 821 (n=7) | 936 (n=5) | 0.106 | 0.371 |
| CXCL13 / BLC | 127 (n=7) | 218 (n=5) | 0.106 | 0.371 |
| IL-17 | 3.49 (n=7) | 2.53 (n=5) | 0.138 | 0.417 |
| CD40 Ligand | 4.71e+04 (n=7) | 5.77e+04 (n=5) | 0.149 | 0.417 |
| IL-7 | 3.75 (n=7) | 2.65 (n=5) | 0.21 | 0.534 |
| TNF-alpha | 103 (n=7) | 148 (n=4) | 0.23 | 0.535 |
| Granzyme B | 28.2 (n=7) | 53.4 (n=5) | 0.268 | 0.535 |
| IP-10 | 675 (n=7) | 1.42e+03 (n=5) | 0.268 | 0.535 |
| GM-CSF | 39.6 (n=7) | 31.3 (n=5) | 0.322 | 0.601 |
| PD-L1 | 1.58e+03 (n=7) | 1.14e+03 (n=5) | 0.343 | 0.601 |
| IL-4 | 0.34 (n=7) | 0.29 (n=5) | 0.365 | 0.601 |
| IL-13 | 71.9 (n=7) | 64.7 (n=5) | 0.492 | 0.707 |
| IL-12 | 16.4 (n=7) | 16.4 (n=5) | 0.497 | 0.707 |
| IL-2 | 2 (n=7) | 4.09 (n=5) | 0.505 | 0.707 |
| CXCL2 / GRO beta | 434 (n=7) | 384 (n=5) | 0.639 | 0.813 |
| IL-8 | 1.12e+03 (n=7) | 1.82e+03 (n=5) | 0.639 | 0.813 |
| IL-6 | 4.59e+03 (n=7) | 4.03e+03 (n=5) | 0.755 | 0.842 |
| IL-15 | 52.7 (n=7) | 58.5 (n=5) | 0.755 | 0.842 |
| PDGF-BB | 11.1 (n=7) | 8.98 (n=5) | 0.755 | 0.842 |
| Eotaxin | 144 (n=7) | 93.5 (n=5) | 0.782 | 0.842 |
| PDGF-AA | 1.07e+03 (n=7) | 963 (n=5) | 0.876 | 0.909 |
| G-CSF | 34.9 (n=7) | 36.1 (n=5) | 0.975 | 0.975 |

**Supplemental Table 8. Luminex assay on fetal plasma.**

| Analyte | AF– Dams | AF+ Dams | P value | FDR |
|---|---|---|---|---|
| TNF-alpha | 0.13 (n=7) | 4.49 (n=5) | 0.00126 | 0.0379 |
| CXCL2 / GRO beta | 838 (n=5) | 263 (n=5) | 0.0794 | 0.758 |
| I-TAC | 1.18 (n=7) | 1.18 (n=5) | 0.152 | 0.758 |
| IFN-beta | 3.25 (n=7) | 3.25 (n=5) | 0.152 | 0.758 |
| IL-21 | 0.37 (n=7) | 0.37 (n=5) | 0.152 | 0.758 |
| CXCL13 / BLC | 11.9 (n=7) | 11.9 (n=5) | 0.152 | 0.758 |
| IL-17 / IL-17A | 0.395 (n=7) | 2.98 (n=5) | 0.187 | 0.801 |
| IL-8 | 772 (n=7) | 450 (n=4) | 0.23 | 0.808 |
| IFN-alpha | 0.01 (n=7) | 1.44 (n=5) | 0.242 | 0.808 |
| GM-CSF | 0.03 (n=7) | 6.49 (n=5) | 0.311 | 0.828 |
| PDGF-BB | 2.59e+03 (n=7) | 977 (n=5) | 0.343 | 0.828 |
| CD40 Ligand | 1.01e+03 (n=7) | 514 (n=5) | 0.404 | 0.828 |
| IL-2 | 3.32 (n=7) | 2.28 (n=5) | 0.407 | 0.828 |
| IL-10 | 6.48 (n=7) | 6.48 (n=5) | 0.417 | 0.828 |
| Granzyme B | 0.45 (n=7) | 0.45 (n=5) | 0.47 | 0.828 |
| G-CSF | 3.52 (n=7) | 3.52 (n=5) | 0.47 | 0.828 |
| MIP-3 alpha | 0.515 (n=7) | 0.515 (n=5) | 0.523 | 0.828 |
| IL-12 p70 | 0.7 (n=7) | 0.7 (n=5) | 0.523 | 0.828 |
| PDGF-AA | 148 (n=7) | 165 (n=4) | 0.527 | 0.828 |
| VEGF | 134 (n=7) | 109 (n=5) | 0.602 | 0.828 |
| Eotaxin | 217 (n=7) | 432 (n=5) | 0.612 | 0.828 |
| IL-15 | 6.33 (n=7) | 5.52 (n=5) | 0.639 | 0.828 |
| IL-7 | 0.48 (n=7) | 0.065 (n=5) | 0.659 | 0.828 |
| PD-L1 | 128 (n=6) | 139 (n=5) | 0.662 | 0.828 |
| IL-6 | 0.875 (n=7) | 1.75 (n=5) | 0.817 | 0.98 |
| IP-10 | 46.1 (n=7) | 45.8 (n=5) | 0.876 | 1.00 |
| BDNF | 3.83e+03 (n=7) | 4.19e+03 (n=4) | 0.927 | 1.00 |
| IL-4 | 0 (n=7) | 0 (n=5) | 1.00 | 1.00 |
| IFN-gamma | 2.15 (n=7) | 2.15 (n=5) | 1.00 | 1.00 |
| IL-13 | 22.4 (n=7) | 22.4 (n=5) | 1.00 | 1.00 |

**Supplemental Table 9. Luminex assay on umbilical cord plasma.**

| Analyte | AF– Dams | AF+ Dams | P value | FDR |
|---|---|---|---|---|
| Granzyme B | 0.45 (n=7) | 4.38 (n=5) | 0.0568 | 0.795 |
| IL-7 | 0.13 (n=7) | 0.065 (n=5) | 0.0808 | 0.795 |
| CXCL2 / GRO beta | 242 (n=6) | 143 (n=5) | 0.0823 | 0.795 |
| G-CSF | 3.52 (n=7) | 3.52 (n=5) | 0.205 | 1.00 |
| VEGF | 142 (n=6) | 120 (n=4) | 0.257 | 1.00 |
| IFN-alpha | 0.01 (n=7) | 1.54 (n=5) | 0.323 | 1.00 |
| IL-6 | 3.66 (n=7) | 20.8 (n=5) | 0.357 | 1.00 |
| I-TAC | 1.18 (n=7) | 1.18 (n=5) | 0.417 | 1.00 |
| IFN-beta | 3.25 (n=7) | 3.25 (n=5) | 0.417 | 1.00 |
| IL-15 | 6.15 (n=7) | 5.79 (n=5) | 0.496 | 1.00 |
| GM-CSF | 0.03 (n=7) | 3.74 (n=5) | 0.52 | 1.00 |
| IL-21 | 0.37 (n=7) | 0.37 (n=5) | 0.523 | 1.00 |
| TNF-alpha | 0.13 (n=7) | 0.13 (n=5) | 0.523 | 1.00 |
| IL-8 | 467 (n=7) | 355 (n=4) | 0.527 | 1.00 |
| CD40 Ligand | 1e+03 (n=7) | 757 (n=5) | 0.639 | 1.00 |
| IP-10 | 41.7 (n=7) | 47.8 (n=5) | 0.639 | 1.00 |
| BDNF | 3.18e+03 (n=7) | 4.15e+03 (n=4) | 0.648 | 1.00 |
| IL-2 | 2.82 (n=7) | 2.03 (n=5) | 0.755 | 1.00 |
| PDGF-AA | 156 (n=7) | 163 (n=4) | 0.788 | 1.00 |
| PD-L1 | 139 (n=6) | 130 (n=5) | 0.792 | 1.00 |
| PDGF-BB | 1.83e+03 (n=7) | 1.78e+03 (n=4) | 0.927 | 1.00 |
| Eotaxin | 277 (n=7) | 279 (n=5) | 0.96 | 1.00 |
| MIP-3 alpha | 0.515 (n=7) | 0.515 (n=5) | 1.00 | 1.00 |
| IL-4 | 0 (n=7) | 0 (n=5) | 1.00 | 1.00 |
| IL-12 p70 | 0.7 (n=7) | 0.7 (n=5) | 1.00 | 1.00 |
| IFN-gamma | 2.15 (n=7) | 2.15 (n=5) | 1.00 | 1.00 |
| IL-10 | 6.48 (n=7) | 6.48 (n=5) | 1.00 | 1.00 |
| IL-13 | 22.4 (n=7) | 22.4 (n=5) | 1.00 | 1.00 |
| IL-17 / IL-17A | 0.395 (n=7) | 0.79 (n=5) | 1.00 | 1.00 |

**Supplemental Table 10. RhCMV primers**

| Name | Sequence | Length |
|---|---|---|
| Rh1-F | CGGGGGGGTGTTTGTGG | 17 |
| Rh1-R | GTAAACCAGGGGCGGTGAAG | 20 |
| Rh2-F | GAACGAGGTGCTTGGATTGTC | 21 |
| Rh2-R | CAGTGGTAACAGATCAGTTTC | 21 |
| Rh3-F | CCTGACTGTAATAGCGGCTG | 20 |
| Rh3-R | CGAATGTCTTTCGGAAACTGG | 21 |
| Rh4-F | GATATTTGTCCGAACACACAAC | 22 |
| Rh4-R | CAGGAGAAAAGGAAATCGCTG | 21 |
| Rh5-F | CTCCACAGCCTCGATGTCTC | 20 |
| Rh5-R | CTCGCCTCATCTTGTGGCTC | 20 |
| Rh6-F | GAATTGATATAAACACTTGACTCC | 24 |
| Rh6-R | CACATAAGCCTGGAAACGTGC | 21 |
| Rh7-F | GCTCGTGCAAAACGTTGCTAG | 21 |
| Rh7-R | GTCGATACCACGTCGTTCTG | 20 |
| Rh8-F | CCATTCACTTGCCTACATGTAG | 22 |
| Rh8-R | CATGTGGTTTGTGTAGTGGTTG | 22 |
| Rh9-F | CCACGATGTCGATAACCAGC | 20 |
| Rh9-R | CTAGTAATCATAGTCCCGTTC | 21 |
| Rh9b-F | CTATGGGGAAAGTATCGTAAAGC | 23 |
| Rh9b-R | CGTAGAATAAGTTGTCGTGGG | 21 |
| Rh10-F | CACTACTGCCCTAACAAATACC | 22 |
| Rh10-R | CTGGGGTGATAGCAAAAACG | 20 |
| Rh11-F | CAACCTAACATGTGACTTTCCAAC | 24 |
| Rh11-R | CCACACTGTTTGTTGGTTTCC | 21 |
| Rh12-F | CATCATGTCCCAGCTGCTATAC | 22 |
| Rh12-R | GTGTTACGGTTCAGTAACTCAC | 22 |
| Rh13-F | TCGCGATTCTGCTGGCGATC | 20 |
| Rh13-R | CTTTACTGTCCTCGCCACATAC | 22 |
| Rh14-F | GGGCCAGATTTCCAAGCATTTC | 22 |
| Rh14-R | GCTGGCTCATGGGGTGATTG | 20 |
| Rh15-F | CCGCAATCTGACATTGCGATC | 21 |
| Rh15-R | ACTTCGTGCGGACCTATCTC | 20 |
| Rh16-F | GAAGACGAAGAAGAGTCAACAG | 22 |
| Rh16-R | CGGTGTACACTTTTCGCTGAG | 21 |
| Rh17-F | CGACTCTCGTCACCCATATTG | 21 |
| Name | Sequence | Length |

| | | |
|---|---|---|
| Rh17-R | CGCTAAGGTAAAAGGCCGTG | 20 |
| Rh18-F | CTGTACGCGTATAATCACAACAC | 23 |
| Rh18-R | GTTTCTGTGGCTGACAATCATG | 22 |
| Rh19-F | GTGATGGCAGTCACTTTACGAG | 22 |
| Rh19-R | GGAGACTCATGGTAGCTCTTTC | 22 |
| Rh20-F | GCTCTCTTACCTATTCAGCTAG | 22 |
| Rh20-R | CGGAGGCCGATGTCGTTGTG | 20 |
| Rh21-F | GCTGAGGCCAGAACTTTTGAG | 21 |
| Rh21-R | CTTGTAGACACGCCTGCTGTC | 21 |
| Rh22-F | GTGCTTTATTACAAATGCAGTGC | 23 |
| Rh22-R | GTGGTGTAATGCCAGTTTCAC | 21 |
| Rh23-F | CCAAGACAAAACACCCCACC | 20 |
| Rh23-R | CAGATTCTGGTACATTCTGCTC | 22 |
| Rh24-F | GAATTCCAACTCACTACCGCTG | 22 |
| Rh24-R | CTCGTGAGGTAGACTGGTTG | 20 |
| Rh25-F | GAGTTAAGTTCTCATACAGCCAC | 23 |
| Rh25-R | CGGCAGTTACTGAAGGGTCG | 20 |
| Rh26-F | CCAATATCTCATACGAAGCGTG | 22 |
| Rh26-R | CAGTTGGAAACACATTGGCTC | 21 |
| Rh27-F | CTATCCAGCGCGCTTCGATC | 20 |
| Rh27-R | CTCTAGCTGTTTGGAAAGTTG | 21 |
| Rh28-F | AGATCTCCATACCGCTAATGC | 21 |
| Rh28-R | GATGCAGAGGTCCTGAATGG | 20 |
| Rh29-F | GCAACTCGACGTGTCCCAGG | 20 |
| Rh29-R | CGACCGCTAGAACCTACCAAG | 21 |
| Rh30-F | CAGCGCCACCGCCAGTC | 17 |
| Rh30-R | TGCTTCAGGCAGATGGTGATG | 21 |
| Rh31-F | TCGACACACGATGCAACCTTTG | 22 |
| Rh31-R | GCTAGTTCTCTCCTCTGTCTTG | 22 |
| Rh32-F | CGATGTGTGGTAAGTTGGATTGC | 23 |
| Rh32-R | GCATTCGCTAGGCGTTTGATG | 21 |
| Rh33-F | CATCGACCTGATCGTAAGTGTG | 22 |
| Rh33-R | GACTTGGTACGGGATCAAGATG | 22 |
| Rh34-F | GCACAGAGAAGCGGGAACC | 19 |
| Rh34-R | GTAGCCAACTCGGTGGGTGTG | 21 |
| Rh35-F | CAGTCGTCGTCAAAGGCATG | 20 |
| Rh35-R | GGTACACGAGTCAGAAGCTCAG | 22 |
| Rh36-F | GAAACCCAGCTCACAGATGAC | 21 |
| **Name** | **Sequence** | **Length** |

| | | |
|---|---|---|
| Rh36-R | GTTCTGGAAAGTTCTGACTCC | 21 |
| Rh37-F | CGCTGACGCTGCGGTTTTTAC | 21 |
| Rh37-R | CTGAGTGTTGTGTGAGTGACAG | 22 |
| Rh38-F | CTGCCTCTCTTTCCCCCCATG | 21 |
| Rh38-R | CATTCGAACGCGAAGAACGTG | 21 |
| Rh39-F | CGTTCGTGGGCGGTCTGAG | 19 |
| Rh39-R | GCACACTCACCAGCCCATC | 19 |
| Rh40-F | GGTCTCCTCCTCATTGAGAC | 20 |
| Rh40-R | AAACAACACAAATCCCGCTTCC | 22 |
| Rh41-F | TTAAAGATGCTCTTCTCCACAAGC | 24 |
| Rh41-R | CGAGACGTATTATACAGGCCTTG | 23 |
| Rh42-F | GTGACTCGATCCTGGATGGTG | 21 |
| Rh42-R | CTTCCTAACGACATGACGCGTC | 22 |
| Rh43-F | ACGAGCTCTTGTCCTGCAATG | 21 |
| Rh43-R | GCGATCTAACATGTTCAACCAC | 22 |
| Rh44-F | TCTGAAACCATGATTACGGAGC | 22 |
| Rh44-R | CTATCGGCCTCCGTGAGATC | 20 |
| Rh45-F | GGCACGTTGGCTGTTTATGG | 20 |
| Rh45-R | ACCAACTGGCCTGTTCCAAC | 20 |
| Rh46-F | GATCAAATTCTCTTCGCCTATG | 22 |
| Rh46-R | CCTTGAACCCATTCCAGAGTC | 21 |
| Rh47-F | CGTCATTCACAACCTAACTACG | 22 |
| Rh47-R | TCGCCAGCCGCATCTCCGTC | 20 |
| Rh48-F | CATGTTTTGGATGGTGGCGTC | 21 |
| Rh48-R | ACATCAACACGACCTACCATG | 21 |
| Rh49-F | GTAGCCGAGGTGTATCCGAG | 20 |
| Rh49-R | CGCTCTGTCGACAGGCCTTG | 20 |
| Rh50-F | AACTCTGTAGGATGTTGCTGC | 21 |
| Rh50-R | GAACCATCCCACATTCACATTG | 22 |
| Rh51-F | GTGTACTGGACGACAAACTTTC | 22 |
| Rh51-R | CTGAATGTGAATGCGCAGGAAC | 22 |
| Rh52-F | GTTGGTGGTAAAGGCCGGG | 19 |
| Rh52-R | GGAGAATTTCTGAGCCACCATG | 22 |
| Rh53 | GAGGTGTATCGCGAAATCAAAG | 22 |
| Rh53-R | TACCGGCTGCAGTTGTCAATC | 21 |
| Rh54-F | CTGTGTCTGTAGTACCGGATG | 21 |
| Rh54-R | GTGCATGAGACAACAGCCTGTG | 22 |
| Rh55-F | CTGGTGTGTATCACGGACTG | 20 |
| **Name** | **Sequence** | **Length** |

| Name | Sequence | Length |
|---|---|---|
| Rh55-R | GAATGTGGTCGCAACTGAACAC | 22 |
| Rh56-F | CCTAACGCACATTTGGTTTCG | 21 |
| Rh56-R | GCTTCCAGGATCTCTAGCTC | 20 |
| Rh57-F | CTCTTATTCGGGTGTTGGCTC | 21 |
| Rh57-R | TTCACACCGACCATGGCATG | 20 |
| Rh58-F | GCTCATTTCGCCGCGGTAC | 19 |
| Rh58-R | CACATCTCCGAAACATCTTCC | 21 |
| Rh59-F | ACGTCTGCAAGTGAACGATCTC | 22 |
| Rh59-R | ACAAAGGATGCACGCCAAAC | 20 |
| Rh60-F | TGAGAGGAGAGAAGTATGAGGTG | 23 |
| Rh60-R | ACTCGATCTACACCACGGAC | 20 |
| Rh61-F | TGGACTACGATGTTTCTCGAG | 21 |
| Rh61-R | GATTCAAGACGCGCGGATCA | 20 |
| Rh62-F | AGCAGGCTAACTCTGTGCAG | 20 |
| Rh62-R | CTACAGTGAGTGTGGAGAC | 19 |
| Rh63-F | GCGCGGCACACATTATCTAC | 20 |
| Rh63-R | CGGACAACCAAGACGTGAAC | 20 |
| Rh64-F | GAGGGTGGGGGATGGATGATC | 21 |
| Rh64-R | GGCTTTTGATTATGATAGTCAG | 22 |
| Rh65-F | CAGACCTACAGTTGTCATGG | 20 |
| Rh65-R | GCAGACTGAGGGAGAGACTG | 20 |
| Rh66-F | TGACAGTCATCTGCAAGTCCTC | 22 |
| Rh66-R | CCATAAGGGGCGGTGCTATTG | 21 |
| Rh67-F | GCAAGCACCGTCACCAATAG | 20 |
| Rh67-R | TATTGATCCATATAGCCAATATCC | 24 |
| Rh68-F | TGGCTATAATCAATACTGGCCCA | 23 |
| Rh68-R | TGAGAGACRGGTARAATTGRCTY | 23 |
| Rh69.1-F | TACCCGTCTCTCAGACCAATT | 21 |
| Rh69.1-R | CTCATGAGGTCTGGGGTTTC | 20 |
| Rh69.2-F | TACCCGTCTCTCAGACCAATT | 21 |
| Rh69.2-R | CTCGTAGAATTGGATTAATA | 20 |
| Rh69.3-F | TGCCCGAGTCTCAGACCAATT | 21 |
| Rh69.3-R | CTCGTAGGATTGGATTAATT | 20 |
| Rh70.1-F | GTCTTTACGTTCGCCACAC | 19 |
| Rh70.1-R | ACGTTATTTATAGCTAATTGTCAT | 24 |
| Rh70.2-F | GATGAGCTTCTTTGCTGAA | 19 |
| Rh70.2-R | AATTGCATATAGTTACGTGCAG | 22 |
| Rh70.3-F | GATGAGTTTCTTTGCTGAA | 19 |

| | | |
|---|---|---|
| Rh70.3-R | AATTGCATATAGTTACGTGCAG | 22 |
| Rh71-F | CGTATGGGTTCAGGTGAATGA | 21 |
| Rh71-R | TTGTGTGCGTGTTCGTAACC | 20 |
| Rh72-F | CCAATCTATCAAACACCGACG | 21 |
| Rh72-R | TCCAAATGGTAACGGAATCTACTGT | 25 |
| Rh73-F | CTCGAGAAGTGACGTTGAAAG | 21 |
| Rh73-R | TCCAAATGGTAACGGAATCTACTGT | 25 |
| Rh74-F | TGCGTACATATTCACGAGTTCCA | 23 |
| Rh74-R | GATCCAGACTGTGCGCCTAAG | 21 |
| Rh75-F | CTCATTGTACTCTGCGCTCC | 20 |
| Rh75-R | GGAGCCCATGTCCTCCATTC | 20 |
| Rh76-F | CCAGGATGGGTGCCGCATC | 19 |
| Rh76-R | AAACGCAAGCAGAGCCAATG | 20 |
| Rh77-F | CAGTGCGGTTTTCACCATGG | 20 |
| Rh77-R | GCTACATCGCCGAGTACCAAC | 21 |
| Rh78-F | AAGAGGCCACATGTCTAAAAC | 21 |
| Rh78-R | CACTGCAACGGAAACACAATG | 21 |
| Rh79-F | CTTTATTTATTTACATGCATAG | 22 |
| Rh79-R | TGCTGTATTTGGCTGCCTCT | 20 |
| Rh80-F | TTAGCTGTCAATCTCTGCAGG | 21 |
| Rh80-R | CAGGTGTAACTCGGGTCGTC | 20 |
| Rh81-F | CGTGTGTGTTAGTCCCCCTC | 20 |
| Rh81-R | ATCGCTTTTGTTGGCCATCG | 20 |
| Rh82-F | ATCTTTCCACCCTTAACATGAC | 22 |
| Rh82-R | GTACTGCTACCACCTGCCTC | 20 |
| Rh83-F | GCAGGTGAAAAGAAGCGCTC | 20 |
| Rh83-R | AGGACCAAGCTATCGCAACC | 20 |
| Rh84-F | CTCAAACATCGGTGGGTGAGG | 21 |
| Rh84-R | TCGGACAGCTAAGCCACTC | 19 |
| Rh85-F | CCCAGATGATAATTCCCATGC | 21 |
| Rh85-R | AACGCTCTAATGAACGAGCCA | 21 |
| Rh86-F | CTTCCAGGGAAACGACCACA | 20 |
| Rh86-R | CCCGGCGTTCGCATAAGATA | 20 |
| Rh87-F | TCTGCTCTTCAAACCCCGAGG | 21 |
| Rh87-R | ACCGTGTCATTTCTCCATGCA | 21 |
| Rh88-F | GACCTCGTCCGACAATTGCA | 20 |
| Rh88-R | CCGAGCTCTCTTCAACCCTG | 20 |
| Rh89-F | GTTCATGCTGTTGCTGCTGC | 20 |
| **Name** | **Sequence** | **Length** |

| | | |
|---|---|---|
| Rh89-R | AGCTCATACCTGAACCGCAC | 20 |
| Rh90-F | GCATAGATGCGGCAGAAACG | 20 |
| Rh90-R | GCACACAGGTACGACGTCAT | 20 |
| Rh91-F | GAGGCGTAACGATAGGATTGG | 21 |
| Rh91-R | ATGCTCCACAGGAAACCGAG | 20 |
| Rh92-F | CTGAAAGCGAAAGGTGTGCC | 20 |
| Rh92-R | GGCTTTAAACATTTTACCAGAAG | 23 |
| Rh93-F | ATCGGAAAGCAATGTACCGC | 20 |
| Rh93-R | GTACATCCACCACTCCGAGG | 20 |
| Rh94-F | GAGCCGGTGTGGATCGAATC | 20 |
| Rh94-R | GTTTTCGGCTGATTCACCCG | 20 |
| Rh95-F | GGACGCTTGACTGACTCGTCA | 21 |
| Rh95-R | GTGCTGCGCATTTCACTTAC | 20 |
| Rh96-F | GCGTGTGTCAAACCAGTTGG | 20 |
| Rh96-R | CAGCGCGYCACCCCGGAGTG | 20 |
| Rh81-F-2 | GTCATTCCCACCGTGTGTG | 19 |
| Rh81-R-2 | CCATTGTCATCGTCGTATGAG | 21 |
| Rh90-F-2 | GAACCATGTAGTTTTCACGAG | 21 |
| Rh90-R-2 | TGACATGAAGGGCAATAAAGC | 21 |
