## Supplementary Figures for "A nonhuman primate model mirrors human congenital cytomegalovirus infection and reveals a spectrum of vertical transmission outcomes"

**Supplemental Figure 1. Viral Loads in Maternal Fluids of Individual Dams.** Plasma, saliva, and urine viral loads of individual dams. Viral loads expressed as mean copies of input RhCMV DNA per mL of plasma, and per ug of input DNA in saliva or urine.

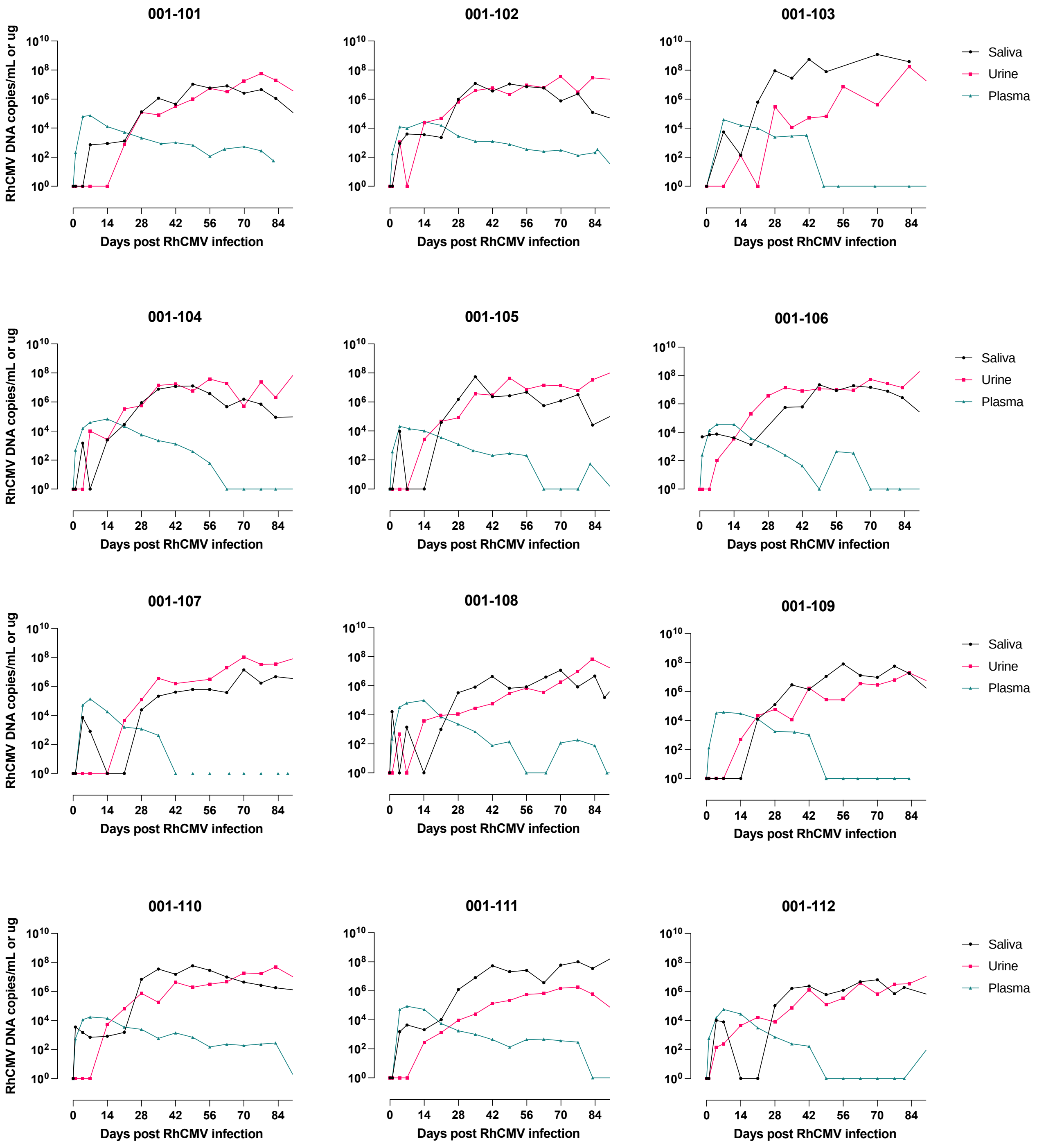

**Supplemental Figure 2. M-F Interface and Fetal Tissue ISH.**

A) In situ hybridization for detection of RhCMV-positive cells in full thickness placenta sections obtained from dam 001-110. Sub-gross view of a full thickness placental section. Annotations were used to delineate the maternal (blue) from fetal (purple) side of the placenta. B) Spatial plots for the visualization of the distribution of RhCMV positive cells within the placenta. Insets: RhCMV+ cells (green) within villi near the fetal (top) and maternal (bottom) interfaces. RNAScope. Green chromogen. No correlation between PCR DNA copies and ISH+ cells in fetal tissues (C) of dam 001-101.

A) B) C)

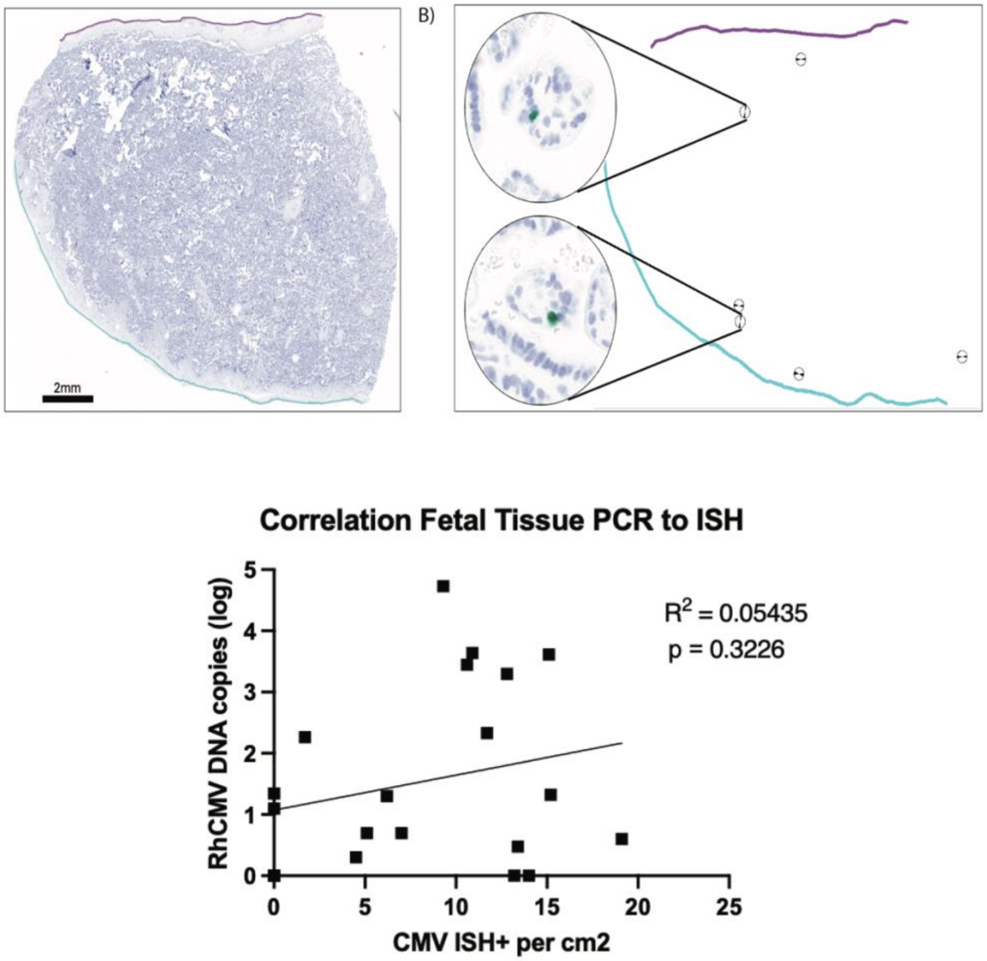

**Supplemental Figure 3. Fetal IgM**

A) Graph demonstrating FL-RhCMV and UCD52 IgG depletion from fetal plasma of all 12 dams. B) IgM ELISA results.

A)

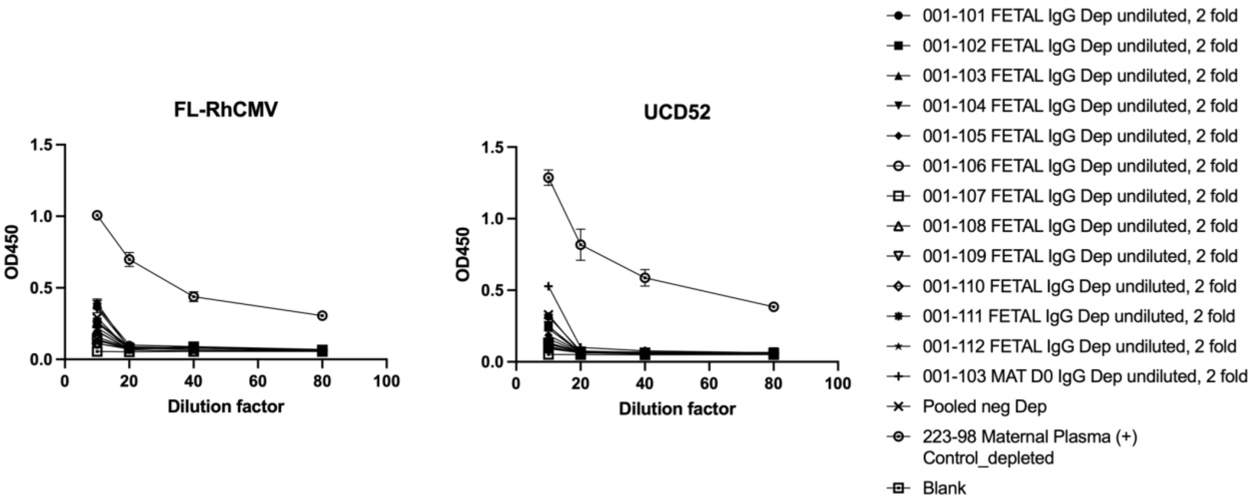


B)

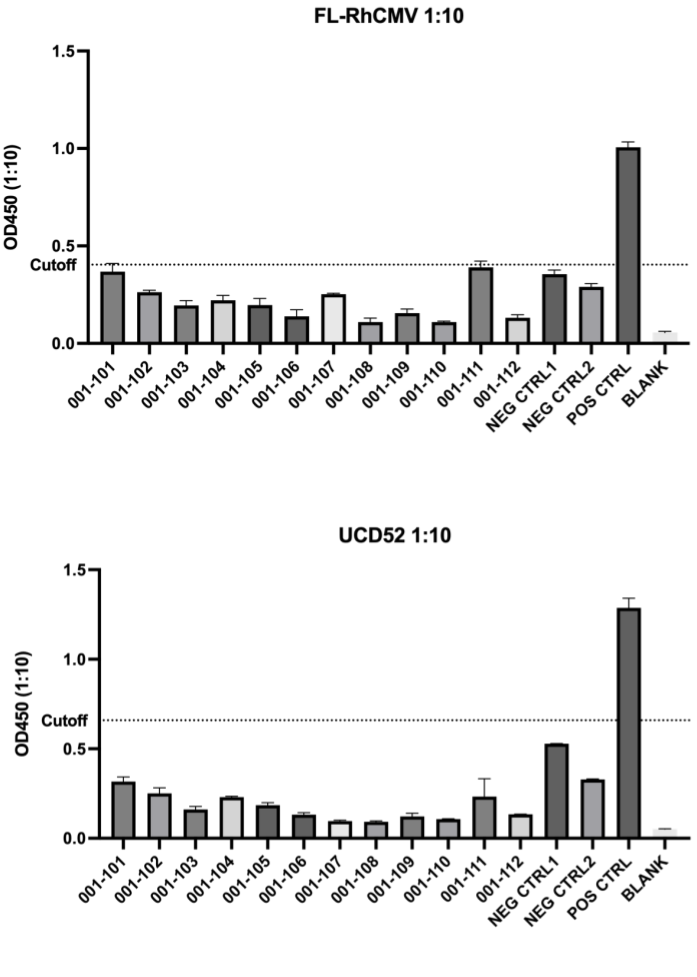

**Supplemental Figure 4. Probability of amniotic fluid transmission and fetal survival.**

A) Kaplan-Meier curve showing probability of amniotic fluid transmission for the study cohort (blue) and the CD4+ T cell-depleted rhesus macaque cohort (n=6) (gray) described in Bialas et al (n=4) and Nelson et al (n=2). B) Kaplan-Meier curve showing probability of fetal survival for the study cohort (blue) and the CD4+ T cell-depleted rhesus macaque cohort (n=6) (gray) described in Bialas et al and Nelson et al.

A)

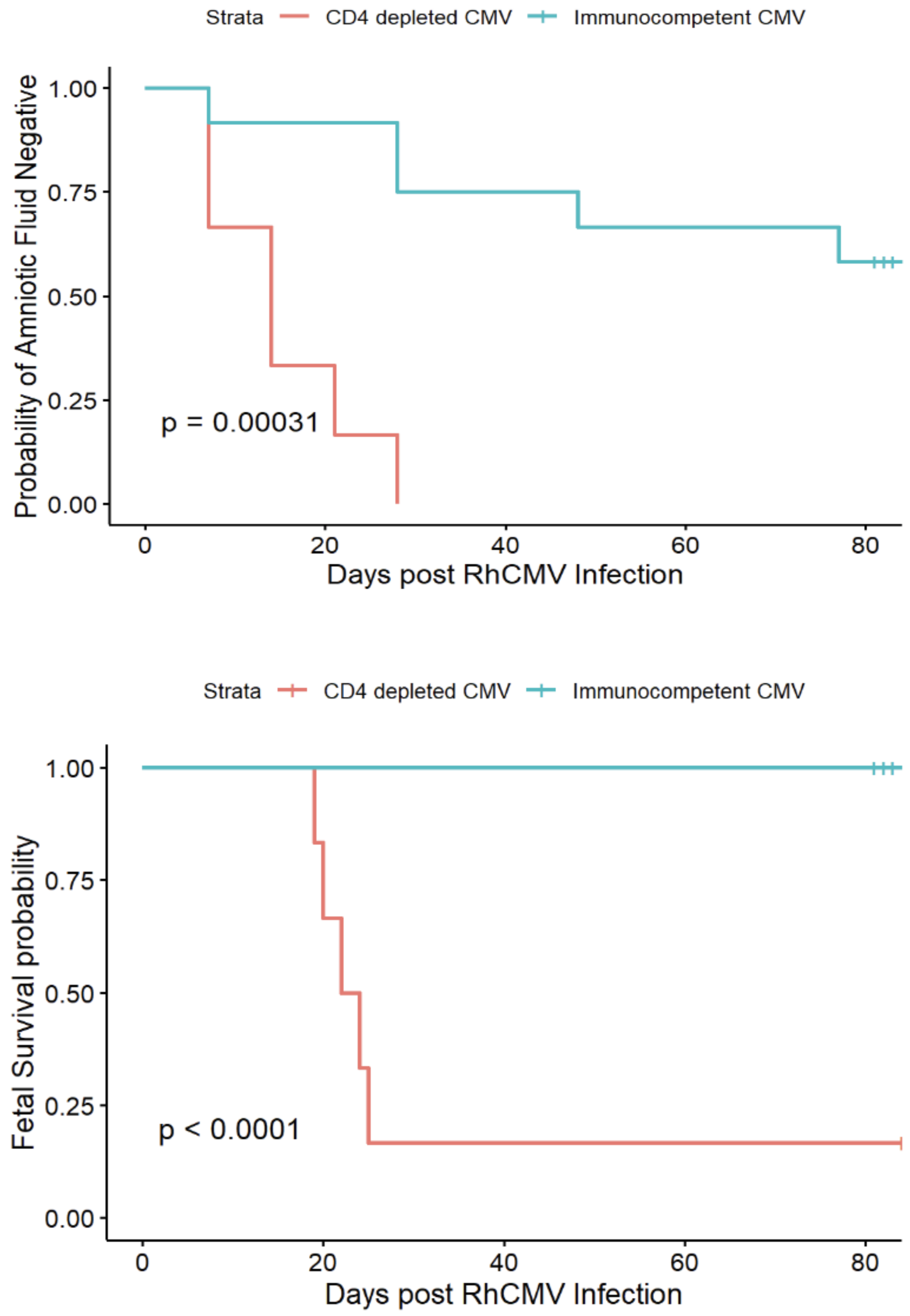

Strata
CD4 depleted CMV
Immunocompetent CMV
Probability of Amniotic Fluid Negative
1.00
0.75
0.50
0.25
0.00
p = 0.00031
0
20
40
60
80
Days post RhCMV Infection
B)
Strata
CD4 depleted CMV
Immunocompetent CMV
Fetal Survival probability
1.00
0.75
0.50
0.25
0.00
p < 0.0001
0
20
40
60
80
Days post RhCMV Infection

**Supplementary Figure 5. Filtering schema for removing strain biased segregating SNVs using FL-RhCMV as the reference.** SNVs positions determined by nucdiff to differ between FL-RhCMV and RhCMV UCD52 (A) are used as the x-axis for the remaining figures. The estimated base fraction at every SNV for each viral inoculum (when mapped to FL-RhCMV) is shown in B and C. Each of the SNV positions were put through three filters (I-L, summarized in D) to identify high-confidence segregating SNVs (E, F). Total read coverage for each inoculum is show in G and H. Right panels show the positions that pass (black) or fail (gray) each of the three filters with threshold shown by the green dashed line: removal of SNVs in hypervariable regions with more than 5 local differences (I), removal of SNVs with fewer than 250 reads when mapped to alternate reference (J), and removal of SNVs with >1% alternate strain frequency in FL-RhCMV (K) or RhCMV UCD52 (L) inoculum. This process was repeated with RhCMV UCD52 as the reference to check for reference strain bias.

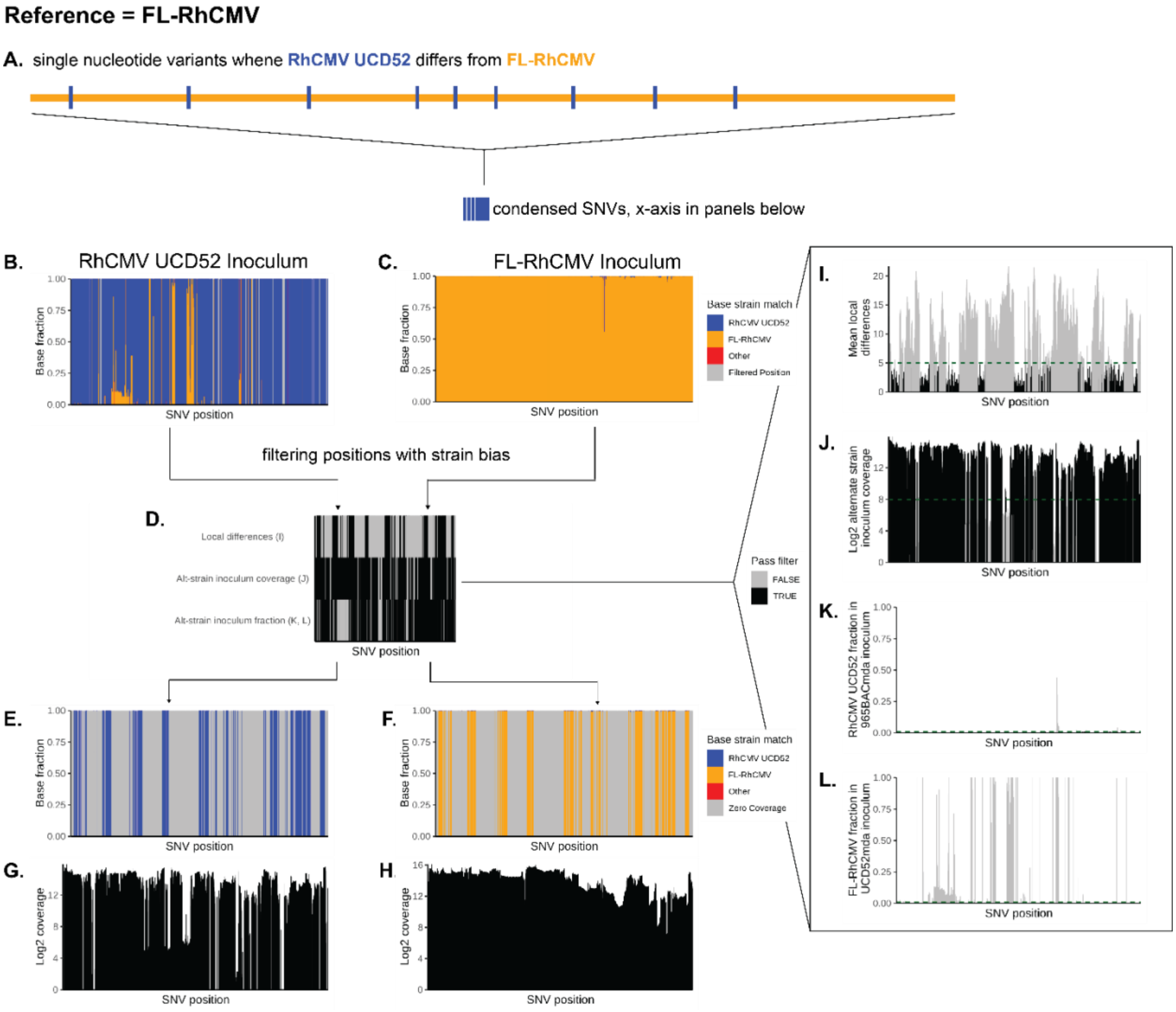

Reference = FL-RhCMV
A. single nucleotide variants whene RhCMV UCD52 differs from FL-RhCMV
condensed SNVs, x-axis in panels below
B. RhCMV UCD52 Inoculum
C. FL-RhCMV Inoculum
Base fraction
SNV position
Base strain match
RhCMV UCD52
FL-RhCMV
Other
Filtered Position
filtering positions with strain bias
D.
Local differences (I)
Alt-strain inoculum coverage (J)
Alt-strain inoculum fraction (K, L)
Pass filter
FALSE
TRUE
E.
F.
Zero Coverage
G.
H.
Log2 coverage
I.
Mean local differences
J.
Log2 alternate strain inoculum coverage
K.
RhCMV UCD52 fraction in 965BACmda inoculum
L.
FL-RhCMV fraction in UCD52mda inoculum

**Supplemental Figure 6. Inferred RhCMV strain frequency in 001-101 with RhCMV UCD52 as mapping reference (top) and sample level strain frequencies (bottom).** A. For each sample (x axis, "P" = plasma, "AF" = amniotic fluid, # = days post infection, M = maternal, F = fetal the inferred strain frequency (blue = RhCMV UCD52, orange = FL-RhCMV) was determined by taking the mean of the percent of reads with the nucleotide corresponding to each strain at the positions where the two inoculating strains differ. Mean indicated by the open circle with error bars corresponding to the standard error. Black dots represent frequencies at a single SNV. In the bottom panels, for each sample, the top panel is the strain frequency calculated with FL-RhCMV as the reference (862 SNVs); the third panel is the strain frequency calculated with UCD52 as the reference (850 SNVs); the second and fourth panels are the read coverage at each position. For each sample, the coloring of each vertical line corresponds to the proportion of reads with the nucleotide that corresponds to either FL-RhCMV (orange) or RhCMV UCD52 (blue). Positions that were excluded from samples due to insufficient coverage (<100 reads) are in grey.

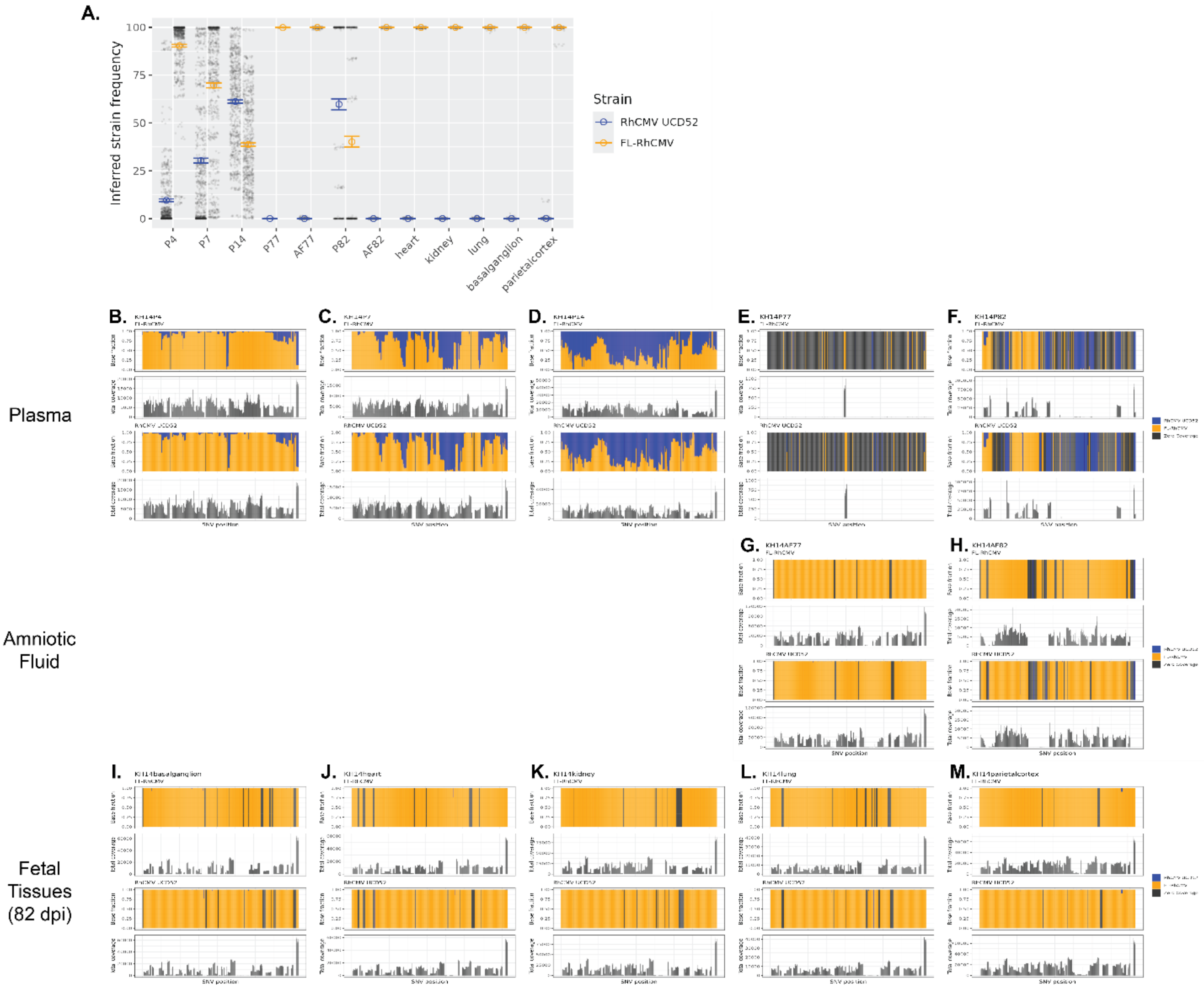
A.
Inferred strain frequency
100
75
50
25
0
P4
P7
P14
P77
AF77
P82
AF82
heart
kidney
lung
basalganglion
parietalcortex
Strain
RhCMV UCD52
FL-RhCMV
B.
C.
D.
E.
F.
G.
H.
I.
J.
K.
L.
M.
Plasma
Amniotic Fluid
Fetal Tissues (82 dpi)

**Supplemental Figure 7. AUC of Viral Loads in Maternal Fluids**

A) AUC of viral loads in plasma, saliva, and urine of AF+ and AF– dams during the first two weeks PI. B) AUC of viral loads in plasma, saliva, and urine of AF+ and AF– dams during the first three weeks PI. C) AUC of viral loads in plasma, saliva, and urine of AF+ and AF– dams during the whole study time course.

A)

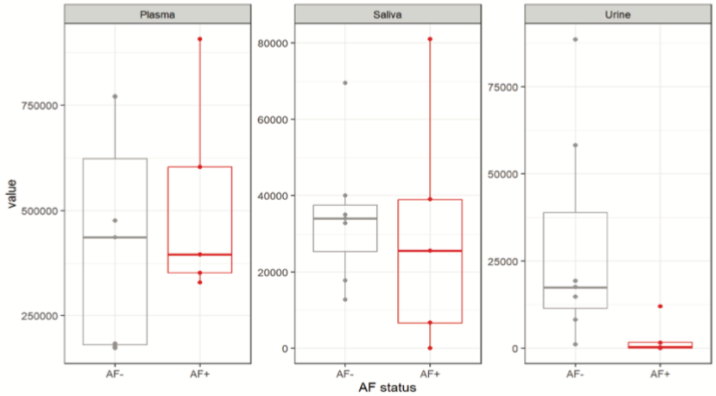


| What | Negative | Positive | P_Wilcox | P_FDR |
|---|---|---|---|---|
| Urine | 1.75e+04 (n=7) | 448 (n=5) | 0.0177 | 0.053 |
| Plasma | 4.36e+05 (n=7) | 3.96e+05 (n=5) | 0.639 | 0.639 |
| Saliva | 3.4e+04 (n=7) | 2.56e+04 (n=5) | 0.639 | 0.639 |

B)

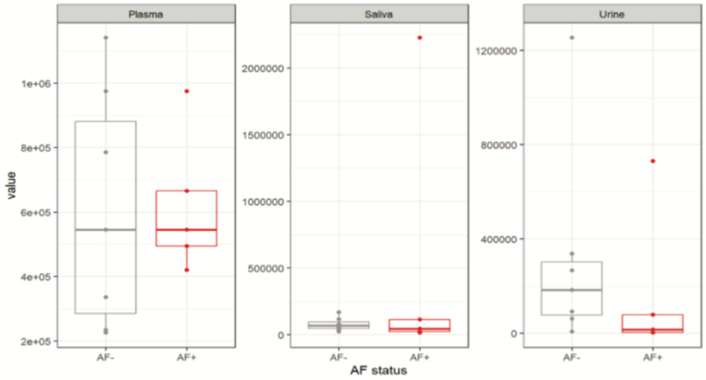


| What | Negative | Positive | P_Wilcox | P_FDR |
|---|---|---|---|---|
| Urine | 1.83e+05 (n=7) | 1.56e+04 (n=5) | 0.202 | 0.606 |
| Saliva | 6.96e+04 (n=7) | 4.49e+04 (n=5) | 0.639 | 0.876 |
| Plasma | 5.45e+05 (n=7) | 5.45e+05 (n=5) | 0.876 | 0.876 |

C)

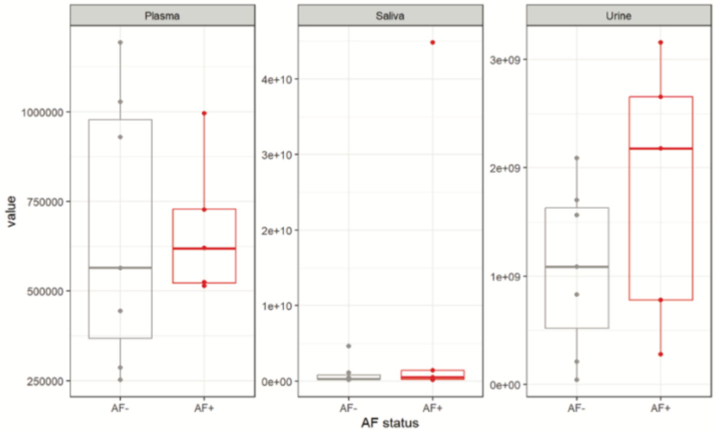


| What | Negative | Positive | P_Wilcox | P_FDR |
|---|---|---|---|---|
| Urine | 1.09e+09 (n=7) | 2.18e+09 (n=5) | 0.268 | 0.803 |
| Saliva | 3.2e+08 (n=7) | 5.31e+08 (n=5) | 0.755 | 0.876 |
| Plasma | 5.66e+05 (n=7) | 6.19e+05 (n=5) | 0.876 | 0.876 |

**Supplemental Figure 8. Chemerin Concentrations in Amniotic Fluid**

Chemerin concentrations in the amniotic fluid taking during the 2nd trimester (between days 14 PI and 28 PI) and at C-section.

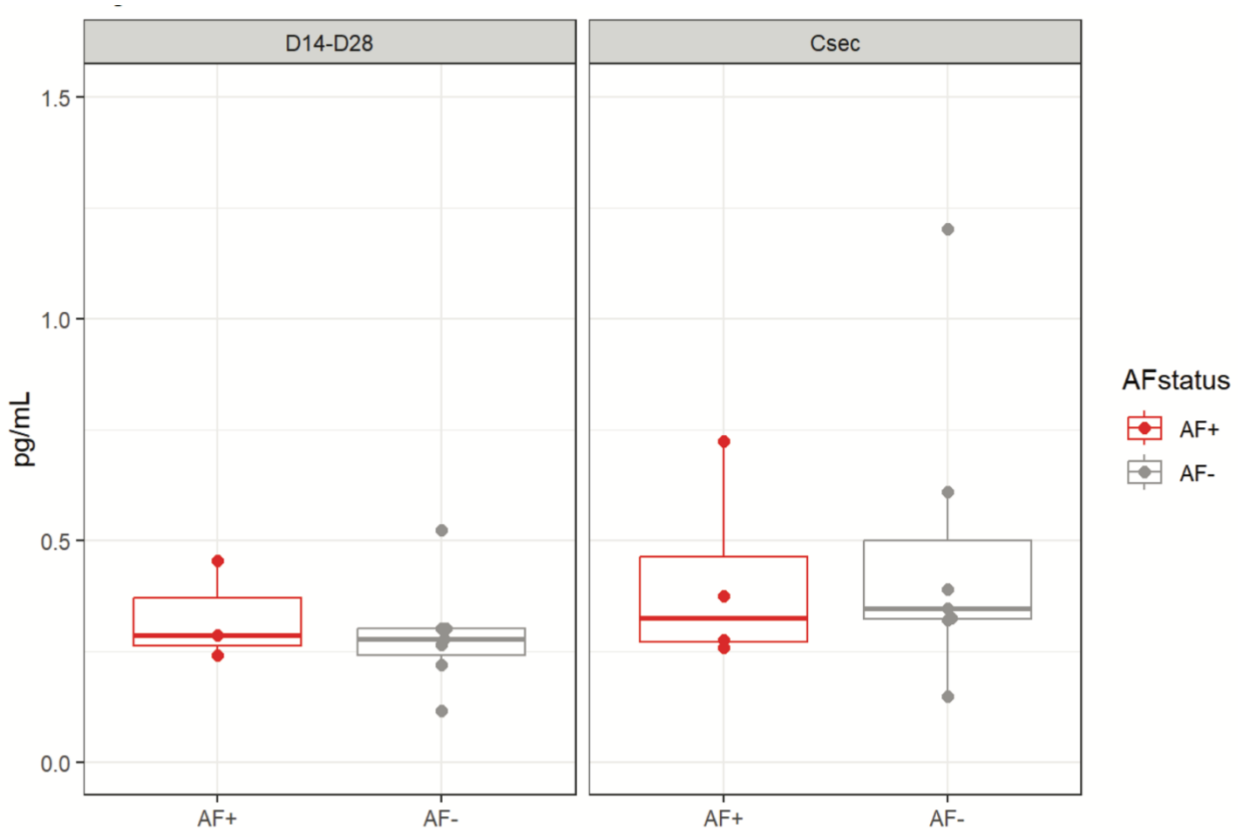
